# Serial colonization shapes genetic variation and defines conservation units in Asian elephants

**DOI:** 10.1101/2023.09.04.556173

**Authors:** Anubhab Khan, Maitreya Sil, Tarsh Thekaekara, Kritika M. Garg, Ishani Sinha, Rupsy Khurana, Raman Sukumar, Uma Ramakrishnan

**Affiliations:** National Centre for Biological Sciences, TIFR, GKVK Campus, Bangalore, India; School of Biodiversity, One Health and Veterinary Medicine, University of Glasgow, Glasgow, UK; National Institute of Science Education and Research, Bhubaneshwar, India; The Shola Trust, Gudalur, India; Centre for Interdisciplinary Archaeological Research, Ashoka University, Sonepat, India; Department of Biology, Ashoka University, Sonepat, India; Centre for Ecological Sciences, Indian Institute of Science, Bangalore, India

**Keywords:** Genomics, genetic load, extinction, megaherbivores, South Asia

## Abstract

Asian elephants (*Elephas maximus*) are the largest extant terrestrial megaherbivores native to Asia, with 60% of their wild population found in India. Despite ecological and cultural importance, their population genetic structure and diversity, demographic history, and ensuing implications for management/conservation remain understudied. We analysed 34 whole genomes (between 11X - 32X) from most known elephant landscapes in India and identified five management/conservation units corresponding to elephants in Northern (Northwestern/Northeastern) India, Central India and three in Southern India. Our genetic data reveal signatures of serial colonisation, and a dilution of genetic diversity from north to south of India. The Northern populations diverged from other populations more than 70,000 years ago, have higher genetic diversity, and low inbreeding/high effective size (Pi = 0.0016±0.0001; F_ROH> 1MB_= 0.09±0.03). Two of three populations in Southern India (South of Palghat Gap: SPG, and South of Shencottah Gap:SSG) have low diversity and are inbred, with very low effective population sizes compared to current census sizes (Pi = 0.0014±0.00009 and 0.0015±0.0001; F_ROH> 1MB_= 0.25±0.09 and 0.17±0.02). Analyses of genetic load reveals purging of potentially high-effect insertion/deletion (indel) deleterious alleles in the Southern populations and potential dilution of all deleterious alleles from north to south in India. However, despite dilution and purging for the damaging mutation load in Southern India, the load that remains is homozygous. High homozygosity of deleterious alleles, coupled with low neutral genetic diversity make these populations (SPG and SSG) high priority for conservation attention. Most surprisingly, our study suggests that patterns of genetic diversity and genetic load can correspond to geographic signatures of serial founding events, even in large, highly mobile, endangered mammals.

## Introduction

Megaherbivores are ecological engineers responsible for maintaining several ecosystem functions (Owen-Smith 1987, Waldram et al. 2008, Coverdale et al. 2016, Berzaghi et al. 2023). They are uniquely responsible for nutrient distribution, seed dispersal and germination, and vegetation/habitat modification (Le Roux et al. 2018, Harich et al. 2016, Sekar et al. 2017). Megaherbivores have been especially impacted negatively in the Anthropocene across their range (Enquist et al. 2020). In Asia, megaherbivores have lost 56% (for gaur, *Bos gaurus*) to near 100% (for Javan rhino, *Rhinoceros sondaicus*) of their historic range (Mahmood et al. 2021).

Conservation genomics suggests that they are more inbred, and lack genetic diversity compared to their small-bodied counterparts (Brüniche-Olsen et al. 2018). While megaherbivore genomes are being extensively studied in African landscapes, such investigations for Asian counterparts have been lagging, leading to challenges in identification of potential genetic threats relevant to their conservation and management.

Asian elephants (*Elephas maximus*) are charismatic megaherbivores distributed across South and Southeast Asia and are culturally important across the globe (Sukumar 2011). They are found in a variety of natural ecosystems from tropical evergreen forests through deciduous forest to grasslands at various elevations. India harbours at least 60% of the population of wild Asian elephants (Sukumar 2011, Menon and Tiwari 2019). Increase in human footprint and land use change over the past two centuries has impacted elephants significantly, resulting in population isolation even at regional scales. Today, Asian elephant habitats are fragmented and interspersed with farmland, human settlement, commercial plantations and linear transport infrastructure throughout their range (Liu et al., 2017; Padalia et al., 2020) leading to extensive and, often, intense human-elephant conflicts (Sukumar, 1989, 2003; Gubbi et al., 2014). We might expect low genetic diversity and high impacts of isolation under such conditions. However, camera trap data (Srinivasaiah et al., 2022; Srinivasaiah et al., 2021) and radiotelemetry studies (Baskaran et al. 1995, Venkataraman et al. 2005, Sukumar 2003, Sukumar et al. 2003) reveal that elephants have annual home ranges of several hundred square kilometres in India, often encompassing dense human habitation. Such movement through human-

dominated areas might offset impacts of fragmentation, and the associated loss of genetic variation and inbreeding (Parida et al. 2022). Clearly, understanding elephant phylogeographic history and current population genetic structuring, coupled with possible effects of recent fragmentation could be useful for elephant conservation and management.

Earlier phylogeographic studies of Asian elephants using mitochondrial DNA point toward a tangled history of demographic change, range expansion/contraction, as well as population admixture possibly mediated by climate change (glacial- interglacial cycles) during the Pleistocene (Fernando et al. 2000, Fleischer at al.

2001, Vidya et al., 2009). These studies showed the presence of two distinct mitochondrial DNA clades (alpha and beta) of Asian elephants, which underwent allopatric separation likely in glacial refugia in Myanmar and Sri Lanka around 1.9 million years ago. Using six (nuclear) microsatellites and a mitochondrial region, Vidya et al. (2005b) reported four Indian elephant population genetic clusters, one each in Northwest-Northeastern India and Central India, and two in Southern India separated by the Palghat Gap in the Western Ghats mountain range, broadly corresponding to their regional population distributions. However, a more recent study by De et al. (2021) suggests only three major genetic clusters corresponding to Northwestern India, Northeastern India, and a combined Southern and Central Indian population (with the Northern (NW&NE) population appearing to be admixed). The patterns of genetic diversity also varied across different ecoregions. Vidya et al. (2005b) showed that the Southern Indian populations harboured lower mitochondrial genetic diversity (haplotype diversity) compared to other populations, while according to De et al. (2021), the Northeastern Indian populations showed low heterozygosity. The only genome-wide study of elephants globally (Palkopoulou et al., 2018), includes poor range-wide sampling of Asian elephants, and suggests old divergences (circa 20-30 kya) between populations in Southern India across the Palghat Gap, as well as between the Northern and the Southern populations. Given the limited genomic data and analyses within Asian elephants, it is difficult to understand demographic history, population genetic structure and, consequently, conservation priorities.

To address these gaps, we used whole genome sequences of wild-caught Asian elephants from all four regions and most landscapes within these in India, encompassing known biogeographic barriers, to assess their population structure, demographic history and genetic variation. Our analyses help infer the predominant factors that have shaped the observed genetic patterns across the subcontinent.

Further, we use this information to propose population management units (MU) and speculate on future genetic issues in elephant conservation.

### Results and Discussion Population structure

We used 28 blood samples collected from wild-born captive elephants of known origin from almost all known elephant landscapes in India (Supplementary Table 1, Parida et al. 2022) and re-sequenced whole genomes (Figure 1). We merged this data with six previously published elephant whole genome sequencing data of which 4 were from India. A set of 2,675,655 SNP markers revealed distinct population structure within elephant landscapes in India. A principal component analysis (PCA), for understanding the clustering of genetic variation, revealed that populations separate from the south to north direction along PC1 axis (Figure 2a). Elephants from Northwestern India and Northeastern India (NW&NE) form a single cluster (Figures 2a, b and c). Populations from Central India (CI) and NW&NE cluster together along the PC axes 1 and 2 but resolve into separate clusters along PC axis 3 and in the ADMIXTURE plot (Figures 2a,b,c,d ) with the most optimum support for five population clusters (Northwest-Northeast, Central and three clusters in Southern India (namely, North of the Palghat Gap (NPG), South of the Palghat Gap (SPG) and South of Shencottah Gap (SSG)), EvalAdmix, Supplementary Figures 1 and 2).

**Figure 1:**
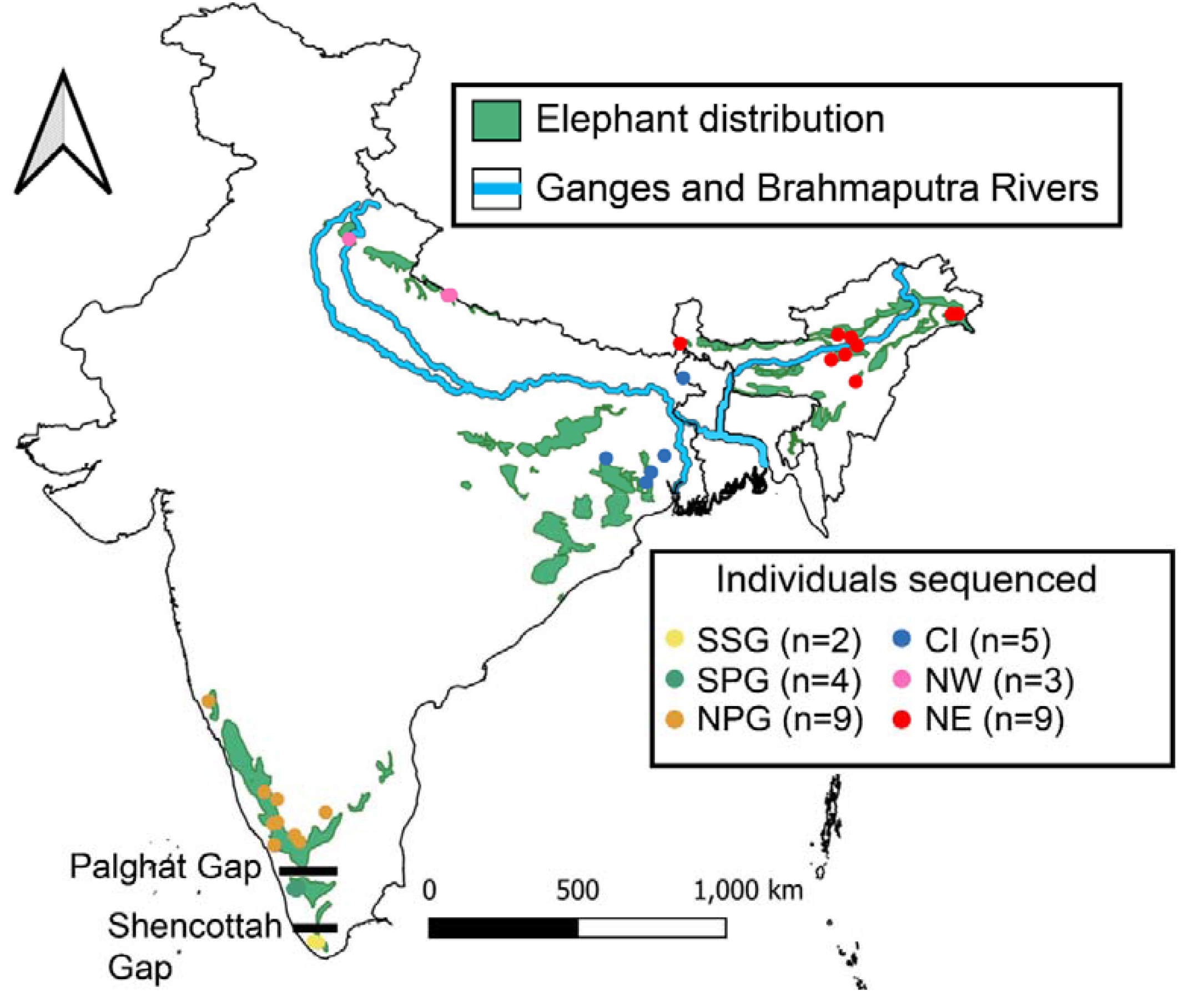
Sampling locations and potential geographic barriers to elephant dispersal. SSG represents south of Shencottah Gap, SPG represents South of Palghat Gap but north of Shencottah Gap, NPG is North of Palghat Gap, CI represents the Central Indian landscape, NW is north-western India, NE is north-eastern India.

**Figure 2:**
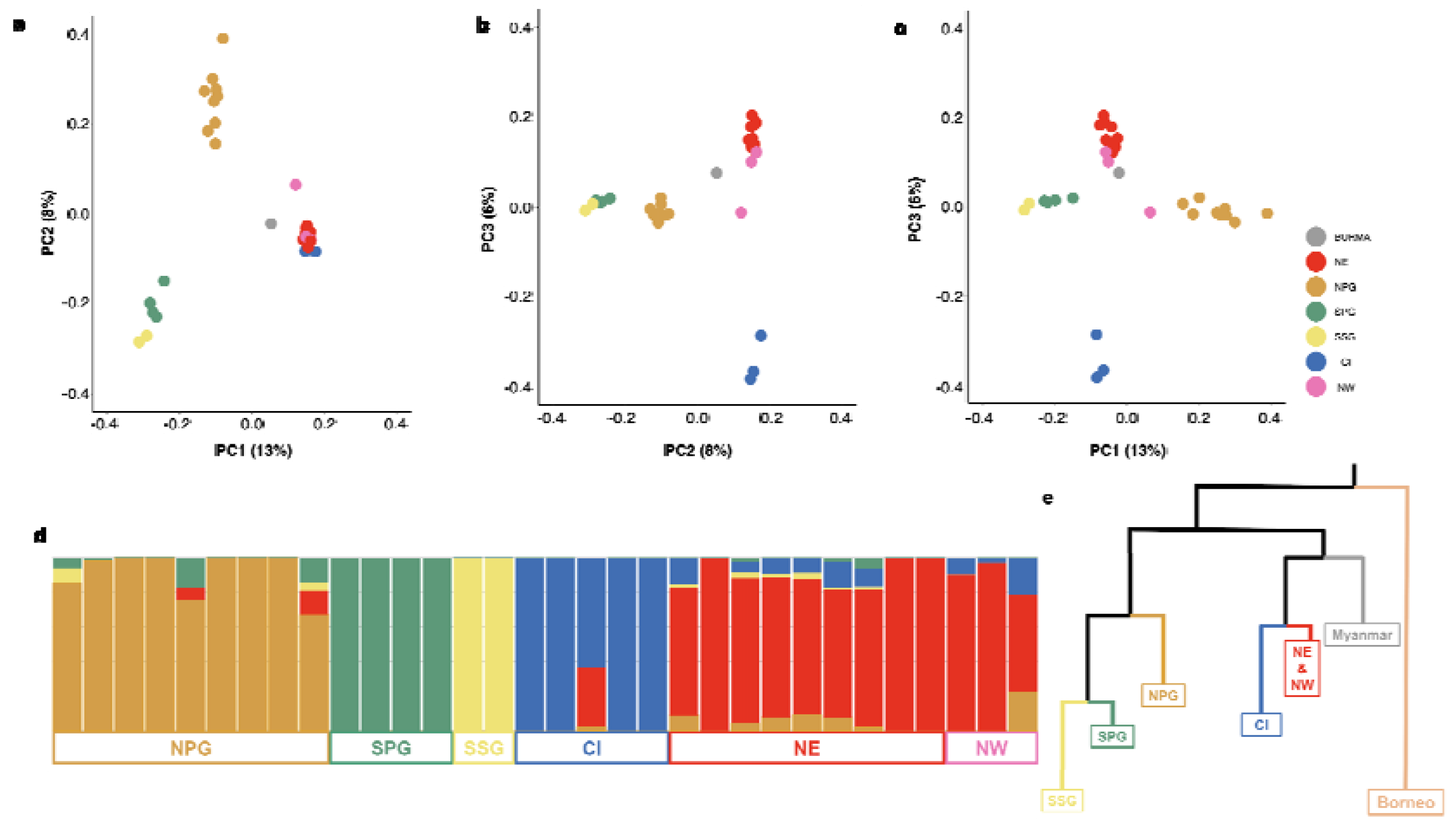
Population structure based on 2.7 million SNPs (a,b,c) PCA, (d) ADMIXTURE plot at K=5, and (e) qpgraph with no admixture events with the branch length qualitatively adjusted to the drift parameter. NW&NE (n=12, sky red) indicates samples from Northeastern India (n=9) and Northern India (n=3), CI (n=5, dark blue) from Central India, NPG (n=9, orange) from North of Palghat Gap, SPG (n=4, green) from South of Palghat Gap but North of Shencottah Gap, and SSG (n=2, yellow) from South of Shencottah Gap.

Northwest and Northeast Indian (NW&NE) populations together, henceforth, are referred to as the Northern Indian population while the population in the south in the Western Ghats (NPG, SPG and SSG) are collectively referred to as the Southern Indian population, unless the subregional populations are being described. The admixture graph that includes a Bornean elephant (ERR2260499) as an outgroup with branch lengths qualitatively adjusted to the drift parameter (Figure 2e,

Supplementary Figure 6) suggests a deep divergence between the Northern and the Southern Indian elephants. Additionally, we observe strong drift parameter values for the two populations South of Palghat Gap potentially indicating low effective population sizes. The phylogenetic (Neighbour Joining or NJ) tree supports the above observed patterns with a minor difference, where the Central population seems to be nested within the Northern population cluster (Supplementary Figure 5).

Our results support previous conclusions that the Northwest and Northeast Indian populations of elephants collectively are different from other Indian populations (Vidya et al. 2005a,b; De et al. 2021) and that the two major northern river systems, namely, the Ganges and the Brahmaputra, have acted as potential barriers to gene flow (Figures 1, 2). Previous studies suggested that the Brahmaputra river acts as an incomplete barrier, with female-led family groups not venturing to cross except perhaps at the upper reaches, but not a barrier to adult male elephants (Vidya et al., 2005b). Largely consistent with these studies, we observe one individual (Rami – female) in the Central Indian samples had admixture with the Northern populations, while another elephant (Bolanath – male) from the Central Indian population dispersed northward across the Ganges river before it was rescued recently from a location close to the Northeastern population (Figure 1, Supplementary Table 1).

Anecdotal information suggests male elephants swim across the Brahmaputra (RS and AK independent personal observations), suggesting it is not a complete barrier to dispersal and movement. Additionally, the Northern Indian elephant samples have ancestries found in all other clusters (Figure 2d), which could be due to incomplete lineage sorting. While geographically proximate, we find that Central Indian populations are genetically distinct as suggested previously (Vidya et al. 2005b).

Consistent with movement ecology inferences (Koirala et al. 2016) the elephants in the Northwestern population (De et al. 2021) are connected to the Northeastern Indian elephants. This is a large landscape running west to east along the Himalayan foothills, though elephant habitat connectivity here is fragile (Koirala et al. 2016) or even completely disrupted at places in recent times (Sukumar 2011, Menon and Tiwari 2019).

Our data and analyses allow identification of a novel genetic cluster in Southern India and suggest three genetically differentiated populations regionally. Along the

Western Ghats in the south, certain breaks or passes divide the elephant population, with the Palghat Gap being the most prominent barrier to elephant dispersal. Further south, the Shencottah Gap also acts as a previously unknown impediment to elephant movement (though anecdotal information suggests that elephants moved across this gap until a few decades ago). Genetic differentiation of SPG and SSG (also see Supplementary Figures 3 and 4) could be due to founder effects and inbreeding combined with recent isolation and small population size (Khan et al.

2022). Gene flow between north and south of the Shencottah Gap may have reduced recently (compared to across the Palghat Gap) largely due to a railway line, a highway and associated development along these transportation infrastructures.

While nested biogeographic implications of these gaps on population structure and phylogeography has been highlighted for smaller species (e.g. montane birds, Robin et al., 2015; bush frogs, Vijaykumar et al., 2017, geckoes, Chaitanya et al., 2019) and some mammals (e.g. Lion Tailed Macaque, Ram et al., 2015), that they result in such deeply divergent lineages in a large, highly mobile mammal such as the elephant is surprising. Alternatively, founding events followed by minimal gene flow might have resulted in the observed patterns of divergence. Additionally, elephant numbers and densities South of the Shencottah Gap have always remained small (on the order of 100 individuals). Such small populations are subject to genetic drift, which could accentuate the observed patterns. Pairwise FST supports these inferences (Supplementary Table 2) and we find no significant gene flow between the clusters based on F3 statistics (Supplementary Table 3). F3 statistics tests whether a target population is an admixture of two test populations. If the F3 value is negative the target population is admixed between the two test populations signifying geneflow between the two populations. For Indian elephants we do not find a significant negative F3 value for any combination of three populations (Supplementary Table 3).

Interestingly, the results obtained from the haplotype network analysis based on the mitogenome paints a slightly different picture (Supplementary Figure 7). Similar to the nuclear genome, the Northwestern population was embedded within the Northeastern population. However, the Central Indian population showed closer affinity to the Southern populations, unlike the results obtained from the nuclear data, and more in line with some previous studies (Vidya et al., 2009, De et al., 2021).

Such discordance between mitochondrial and nuclear datasets in phylogeographic interpretations has been reported previously too (Toews and Brelsford 2012). Since elephants form matrilineal herds, individuals in a herd should share their mitochondrial haplotype. It is possible that herds of matrilineally-related elephants from the Northern population colonized Central India (CI) and North of Palghat Gap, and subsequently founded the other southern populations, leading to the observed patterns.

### Demographic History

We investigated recent demographic history for clusters with more than nine samples (Northern and NPG) using GONE (Santiago et al. 2020) and show both these populations have undergone a recent bottleneck around 1500-1000 years ago (Figure 3a). Since the taming of elephants in the subcontinent during Harappan times, at least four millennia ago, there has been regular exploitation of wild individuals for military and domestic use (Sukumar 2011). We do not have estimates of the actual extent of exploitation (in terms of annual offtakes) but can only qualitatively infer the levels of exploitation from the stocks of elephants in captivity. Historical accounts suggest that the armies of ancient kingdoms and republics in the north (the Gangetic basin) maintained several thousand captive elephants from as early as the 3^rd^ century BCE until late Mughal times in the early 17^th^ century CE, suggestive of overexploitation of wild populations for nearly two centuries, until the invention and use of gunpowder in warfare rendered the war elephant irrelevant (Sukumar 2011). This is the first report of a bottleneck in Asian elephant populations in recent historical times, potentially mediated by human exploitation. Our results also suggest that both populations (Northern and NPG) may have started recovering from the bottlenecks around 300-500 years ago. However, the modelled “recovery” could also be an artefact due to our under-sampling, or population structure. The recent recovery-like pattern may be observed due to recent and fine-scale population structure in the Northern cluster (Sukumar 2011, Menon and Tiwari 2019), which would reduce shared linkage disequilibrium (LD) blocks between populations, giving an illusion of population expansion (Novo et al., 2023). Often, population structure can dictate the models of demographic history (Mazet et al. 2016).

**Figure 3:**
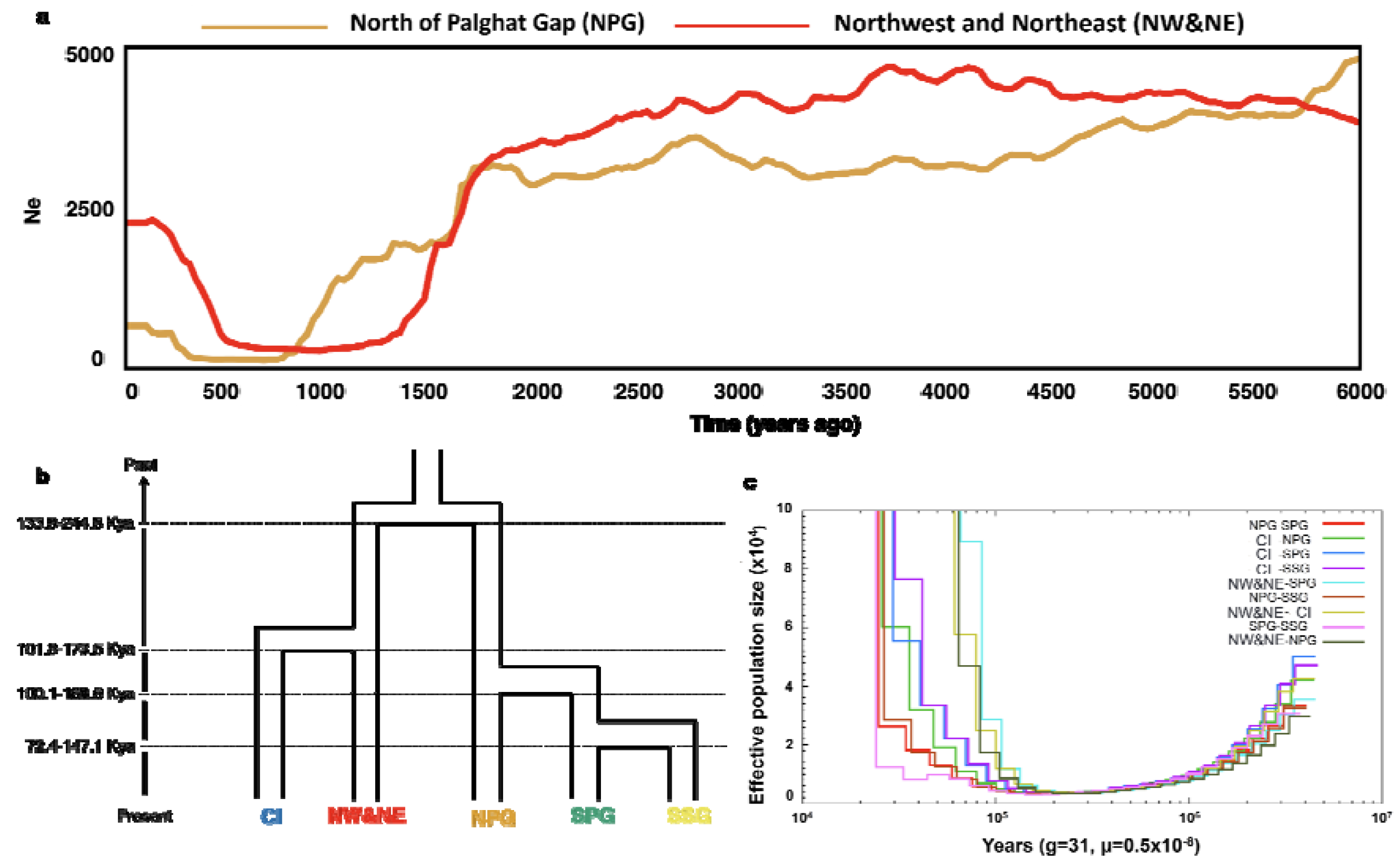
Demographic history (a) recent demographic history within in the last 100 generations inferred from GONE (Santiago et al, 2020) (b) most likely demographic model obtained from fastsimcoal2 (Excoffier et al. 2021) (c) population divergence estimated from hPSMC (Cahill et al. 2016). The point where population sizes start increasing exponentially in hPSMC plot is the time of population divergence. The colours used to represent the populations remain the same as earlier, Northwest and Northeast (sky blue), Central India (dark blue), North of Palghat Gap (orange), South of Palghat Gap (green) and South of Shencottah Gap (yellow).

Additionally, we find signatures of population bottlenecks around 100,000 years ago (Supplementary Figure 11). However, further investigation with hPSMC (Cahill et al. 2016) and fastsimcoal 2 (Excoffier et al. 2021) suggests that this coincides with populations differentiating from each other (Figure 3 b,c). hPSMC results suggest that the Northern elephant population separated from all other populations about 70,000-100,000 years ago while *fastsimcoal* suggests it split from the Southern Indian populations 134-245 Kya and from the Central Indian (CI) cluster 102-174 Kya. The hPSMC results further suggest that the Central Indian elephants diverged from the rest around 50,000-80,000 years ago, while the three Southern Indian populations diverged from each other only around 20,000-30,000 years ago.

However, *fastsimcoal2* supports a model with Southern Indian populations splitting from each other between 72-100 Kya. The serial reduction of historic effective population size from north to south indicates a potential serial founding of populations of elephants in India along this direction (Slatkin & Excoffier 2012).

Overall, our results emphasise the antiquity of the Northern populations of elephants consistent with Vidya et al. (2005b, 2009). Palkopoulou et al. (2018) had suggested a more recent divergence which could be due to the differences in the mutation rates used for making inferences (4.06e-8 vs 5.3e-9; Prado et al. 2023). Vidya et al. (2009) made inferences on phylogeography based solely on mitochondrial DNA sequences, which are expected to provide older estimates of divergence times even up to an order of magnitude greater (Zheng et al. 2011). Discordance between mitochondrial and nuclear genome estimates of phylogeography has been attributed to several reasons include sex-biased dispersal, faster sorting rate of mtDNA than nuDNA and differences in the geographical range or numbers of hybridizing populations (Peters et al. 2012, Toews and Brelsford 2012); thus, inferences of phylogeny and divergence times from mtDNA alone are best avoided (Weins et al. 2010). However, consistent with Palkopoulou et al. (2018) who used whole genomes, we find that the NPG and SPG populations in the south diverged from each other only around 20,000 years ago, or the time of the Last Glacial Maximum when southern India, in particular, the Western Ghats, was more arid (Sukumar et al. 1993, Rajagopalan et al. 1997). These findings further support the recognition of five elephant management units in India, emphasising their antiquity and unique evolutionary histories.

### Genetic Diversity

To measure how the management units compare with each other, we estimated the genetic diversity as the number of average pairwise differences in sequences (pi) within each cluster. We find that the clusters that are from Southern India (SPG, SSG and NPG) have lower average nucleotide diversity than the Northern (NW&NE) Indian and Central Indian (CI) clusters. Interestingly, there is no visible difference in nucleotide diversity within the Southern and between the Northern and Central clusters (Figure 4a) (Supplementary Table 4). This suggests that all the clusters within Southern India have a similar number of haplotypes and that the Central and Northern cluster have a similar number of haplotypes. Expectedly, the only significant statistical differences were observed between Northern and the Southern clusters, and Central and the Southern clusters, respectively (Supplementary Table 5). About 75% of the discovered SNPs are shared, or present in all populations (Supplementary Table 6). However, the number of heterozygous sites encountered per Mb is the highest for the Northern population while it is lowest for the southernmost population (SSG) (Figure 4b). We find no difference in heterozygous SNV per Mb for NPG and CI (Supplementary Table 5). Overall, our results suggest that most populations of elephants have similar nucleotide diversity but, in the population South of the Shencottah Gap (SSG), similar nucleotides are often paired together, indicating lower effective population size, while in the populations in Northern India most of the nucleotides pair with a different one, indicative of higher effective size. These patterns are consistent with signatures of serial founding, where it is predicted that the heterozygosity of each successive population decreases in comparison to the parent population and is proportional to the effective population size of the founders (Slatkin & Excoffier 2012).

**Figure 4:**
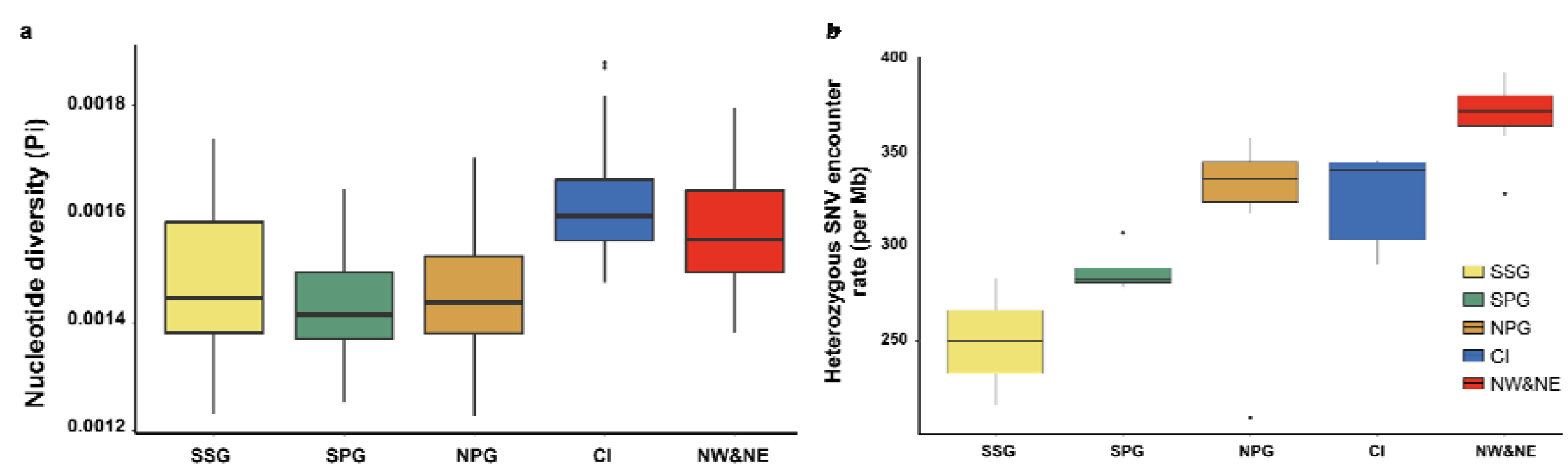
Genetic diversity revealed as (a) boxplots o f pairwise nucleotide differences per site (pi) in the populations (error bars represent variance in mean pi/site across scaffolds), and (b) heterozygous SNV encounter rate per Mb of the genome. The populations are represented, as earlier, the different colours include Northwest and Northeast (red), Central India (dark blue), North of Palghat Gap (orange), South of Palghat Gap (green) and South of Shencottah Gap (yellow).

Genetic variation is related to effective population size which, in turn, is dependent not only on the census population size of a species but other variables such as the adult sex ratio (Frankel and Soulé 1981). Although we cannot speculate on the historical population sizes of Asian elephants across the various regions we have investigated, a cursory look at the recent population sizes of the five management/conservation units we have inferred from the genomic data shows that census population size is not indicative of the observed genetic variation (pi and heterozygosity) (Supplementary Table 7). For instance, the genetic variation of the Northeastern (NE) population is much higher than that of the Southern population to the North of the Palghat Gap (NPG). Both these clusters have similar census population size on the order c. 10,000 individuals. Central Indian elephants have higher variation than Southern populations even though the former has distinctly lower census population size of c.3000 elephants (Supplementary Table 7). This might be indicative of recent bottlenecks in Central India, as heterozygosity decays slowly, while the Southern populations, especially those South of the Palghat Gap may have had historically smaller populations. The highly female-biased sex ratios in Southern populations, especially those to the South of the Palghat Gap, in recent historical times (1970s-1990s) from selective poaching of male elephants for ivory (Sukumar 1989, Ramakrishnan et al. 1998), could have decreased the effective population size (compared to census size). Elephants to the South of Shencottah Gap were also connected to the SPG population through movement of males until the early 1980s according to anecdotal evidence (Gangadharan et al. 2011, Johnsingh et al. 2009). Overall, our results are thus also consistent with a serial dilution of variation that could be the result of sequential colonization (Hellenthal et al., 2008; Pierce et al., 2014; Pless et al., 2022) from north to south: from Northern to Central and Northern to Southern, and then from North of Palghat Gap to South of Shencottah Gap, in that order.

We further test this by estimating FROH, the proportion of the genome in homozygous stretches, with longer stretches indicative of recent bottlenecks and/or inbreeding (Ceballos et al. 2018). We observe that individuals from South of the Shencottah Gap (SSG) have high average FROH>0.1Mb of 0.4 (40% of the genome is in homozygous stretches) while individuals from Northern India have the least average FROH>0.1Mb of 0.2 (20% of the genome is in homozygous stretches, Figure 5a). There is no significant difference between individuals from SPG, NPG and CI populations (average FROH>0.1Mb is 0.25). Longer homozygous stretches are indicative of recent inbreeding (Curik et al., 2014; Sumreddee, 2021). ROH longer than 1Mb and 10Mb shows few differences between populations, although the Northern Indian population has the least recent inbreeding. Additionally, the SPG and Northern populations have lower variance in FROH compared to other populations, potentially indicating gene flow and connectivity within these large landscapes, which will result in lower

**Figure 5:**
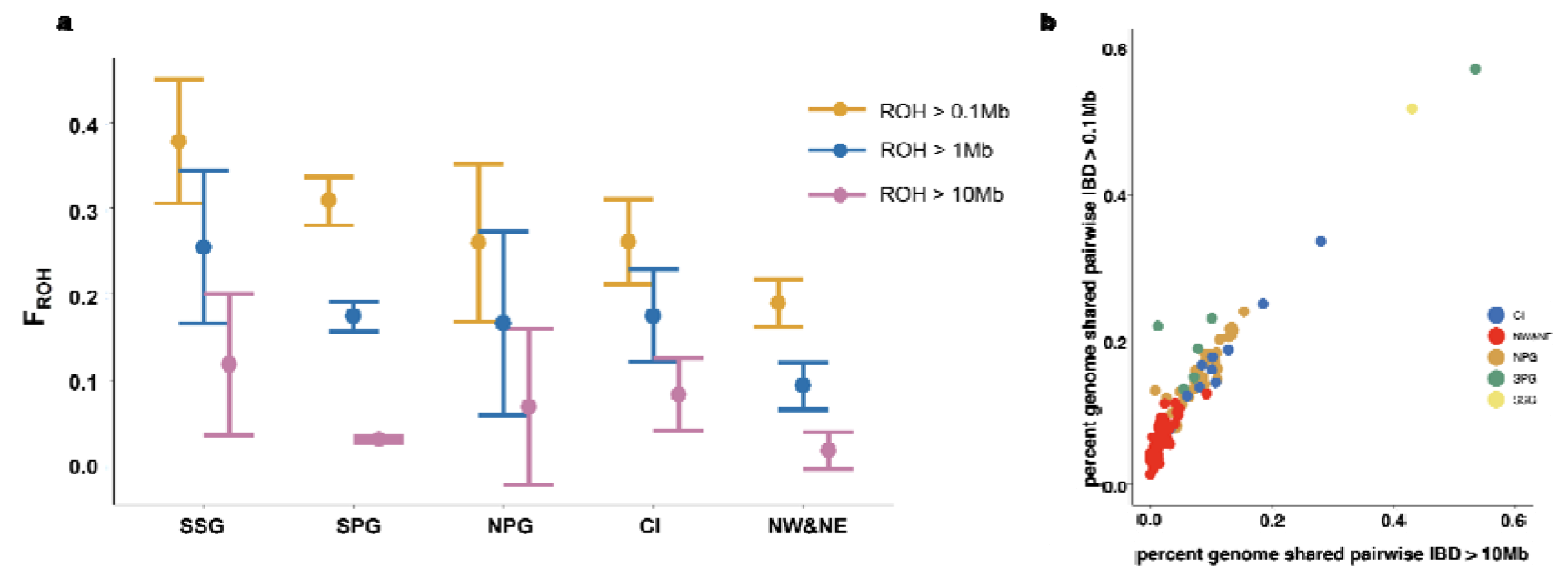
Present and potential future inbreeding (a) inbreeding measured as FROH of individuals in each population based on ROH stretches longer than 0.1 Mb (mustard), 1 Mb (blue) and 10 Mb (pink) of genome and (b) identical by descent (IBD) stretches of the genome longer than 0.1Mb vs 10Mb shared between pairs of individuals in each population. The populations are represented, as earlier, the different colours include Northwest and Northeast (red), Central India (dark blue), North of Palghat Gap (orange), South of Palghat Gap (green) and South of Shencottah Gap (yellow).

variance in parental relatedness. The results suggest that despite inbreeding avoidance in elephants (Sukumar 2003), founder effects and drift led to highly inbred individuals.

While the FROH analyses suggest individual inbreeding, a high average FROH in a population need not result in inbreeding depression, as the homozygous segments may differ between individuals within a population. To measure the consequences of mating between individuals within each population, we measured the percent genome that is identical by descent (IBD) shared between pairs of individuals in each population (Figure 5b). Elephants in the Northern Indian cluster (NW&NE) share very few IBD stretches of the genome (on an average 1.6% in more than 10Mb long and 6.6% in more than 0.1Mb long IBD stretches) with each other, while the two individuals we had from South of Shencottah Gap (SSG) shared about 43% of their genome in stretches longer than 10Mb and about 52% of their genome in stretches longer than 0.1Mb. Apart from the one outlier pair in the SPG sample set, we observe negligible differences in the SPG, NPG and CI populations with regards to the proportion IBD stretches of genome shared between pairs of individuals. On average these three clusters share more than 10% of their genome in IBD longer than 10Mb (NPG: 8%, SPG: 14%, CI:12%) and 19% of their genome in IBD longer than 0.1Mb (NPG: 15%, SPG: 25%, CI:17%). These results indicate inbreeding depression in the future in SSG, SPG, CI populations, due to inbreeding in upcoming generations. Since the present individuals already share large IBD stretches of the genome, impacts on fitness will be maximal, especially if these individuals have higher numbers of deleterious alleles. However, since there are a few outlier pairs with lower amounts of shared genomic IBD stretches (Figure 5b), there is hope that maintaining gene flow within these population clusters could sustain extant genetic variation in these populations into the future.

### Genetic load

We attempted to understand whether individuals, from certain populations, that have high homozygosity could suffer fitness effects due to inbreeding depression by estimating genetic load, or putative damaging mutations. This depends on the number of derived deleterious mutations harboured across the genome, how many are homozygous, and the putative magnitude of their fitness effects (indel Loss of function > Loss of function > missense mutation) (Barton & Zeng 2018; Khan et al. 2021). We normalised this by derived SNPs in neutral/synonymous/intergenic regions, to account for possible missingness/uneven coverage across genomes. As expected, populations with lowest genetic variation (SSG) had fewer LOF mutations (6a, x axis). This could be because of the serial founding demographic history from north to south (see Figure 1e). However, LOF mutations in SSG were not entirely a subset of those in the Northern population, potentially supportive of the emergence of *de-novo* mutations in the SSG population along with differential influence of drift and selection due to the long history of isolation between the two populations. The SSG population has the highest homozygosity for the LOF mutations. Interestingly, SPG also had fewer homozygous LOF mutations (Figure 6a), potentially due to higher effective population size and *de-novo* mutation accumulation due to historical isolation.

**Figure 6:**
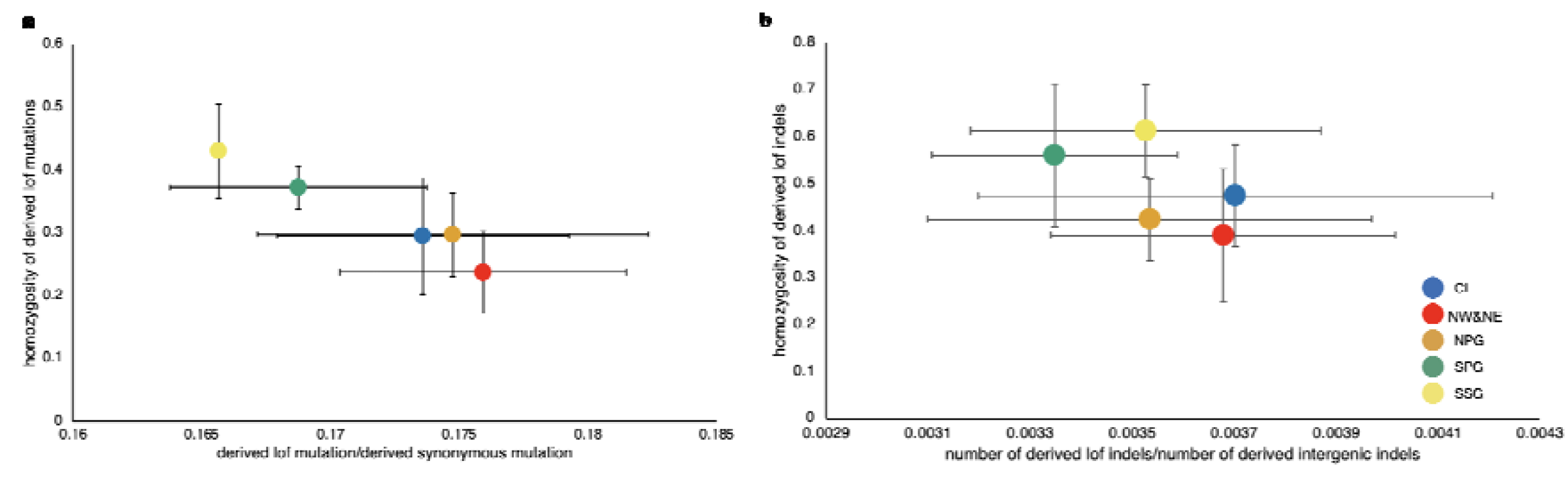
Mutation load measured as a function of homozygosity vs number of derived (a) loss-of-function (LoF) mutations and (b) indel mutations. The number of derived deleterious alleles/number of derived neutral alleles is a proxy for number of deleterious alleles. The error bars indicate standard deviations. The populations are represented, as earlier, the different colours include Northwest and Northeast (red), Central (dark blue), North of Palghat Gap (orange), South of Palghat Gap (green) and South of Shencottah Gap (yellow).

To better understand whether elephant populations such as SSG are endangered by imminent inbreeding depression, we compared number and homozygosity of indel LOFs between all five populations. The total indel LOFs did not vary considerably between populations (x axis, Figure 6b). The observation that all populations have a similar indel LOF load (compared to LOF load, Figure 6a, x axis) suggests that they are under strong purifying selection, and that these putative high negative effect mutations could have already been purged from these populations. In contrast, the homozygous indel LOFs remain higher in SSG and SPG. This suggests that despite purging, elephants to South of the Palghat Gap could experience negative fitness consequences of indel LOFs first. This population may be threatened with inbreeding depression, or lower fitness of individuals, ensuing negative feedback on population growth rates, and a potential for further increase in homozygosity due to inbreeding in the future generations (Robinson et al. 2023). However, the Asian elephant in Borneo, despite its extremely narrow genetic base (Goosens et al. 2016), possibly the outcome of a severe bottleneck during colonization of the island in the last glacial maximum (Sharma et al. 2018), has recovered and maintains a population of about 2000 individuals. Also, African elephant populations (such as at Addo National Park, South Africa) grew rapidly with no apparent deleterious effects after a severe bottleneck about a century ago (Whitehouse and Hall-Martin 2000, Whitehouse and Harley 2001). Genetically small populations are expected to be especially efficient at purging strongly deleterious alleles (Kyriazis et al. 2023, Khan et al. 2021), this does seem to be happening in the isolated population to South of the Shencottah Gap leading to fewer number of deleterious alleles there. But the question that remains is whether these endangered populations with long generation times and low population growth rates can tolerate high genetic load.

Southern populations (NPG, SPG and SSG) have fewer derived missense mutations than the Central (CI) and the Northern populations (Supplementary Figure 8) but the populations South of Palghat Gap (SPG and SSG) have higher homozygous missense mutation load than the NPG, CI and Northern populations, again potentially due to drift and inbreeding. The fewer number of mildly deleterious missense alleles in the Southern Indian population to the North of Palghat Gap (NPG) compared to CI and Northern is counterintuitive, as current literature suggests that larger populations (Supplementary Table 7) host high numbers of mildly deleterious alleles (Kyriazis et al. 2023, Khan et al. 2021). We suggest serial dilution of missense variants combined with a long isolation from the larger and connected Northern Indian population may have led to restricted immigration of mild impact deleterious alleles in the NPG population while the LOF are purged out equally well in the large NPG, CI and Northern populations. Additionally, the low homozygosity in the NPG, CI and Northern populations indicates a potential for inbreeding depression (Robinson et al. 2023). Overall, we observe that the populations in Northern India have high numbers of deleterious alleles but the effect of these alleles are masked due to heterozygosity. Although, the Southern populations have fewer deleterious alleles, they have a high chance of expressing them due to high homozygosity. For example, in the southernmost population (SSG) more than 40% of the LOF mutations are homozygous while in the northernmost population (NW&NE) about 20% of the LOF mutations are homozygous. Thus, serial founding in elephants seems to increase the realized genetic load in every successive founding event.

While the exact fitness effects of mutations, and the distribution of fitness effects is unknown in most wild species, we tried to better understand the possible fitness effects of load. The LOF mutations were annotated using the FUSIL database (Cacheiro et al. 2020). The database creates knock-out mutations of specific genes in mice and evaluates if specific knock-out mutations are lethal, sub-viable (only 12.5% individuals survive) or harmless. Among the genes harbouring LOF mutations in Indian elephants, 892 were represented in the FUSIL database. Out of these 892 genes, 188 LOF mutation were found to be homozygous lethal in mice and 112 mutations were found to lead to a sub-viable phenotype in mice. Thus, about 34% of the LOF mutations, when homozygous, produce a deleterious phenotype in mice.

This highlights the risks the elephants face in near future. While the mutations causing lethal phenotypes are expected to be purged out faster, the ones with sub- viable phenotype may pose the real threat and lead to a slow decline in elephant populations.

Functionally, the derived missense mutations are related to sensory perception, detection of chemical stimulus and especially olfaction (Supplementary Figure 9). However, there is an abundance of olfactory genes in the elephant genome (Reddy et al. 2015) and thus there are more opportunities for mutations. Additionally, several olfactory genes can be pseudogenes. However, since we also polarized the alleles with the Bornean elephant genome, the missense mutations can be confidently classified to be recently derived and maybe potentially deleterious. The loss-of-function mutations mostly affect protein, ions and nucleic acid binding abilities along with transferases and transporters (Supplementary Figure 10). Although inbred individuals are more homozygous for these mutations and maybe expected to be severely affected, it is possible that sociality and herd living allows reduced sensory abilities in individuals. However, this needs further investigation.

### Conservation implications

Overall, our results suggest five management units for Asian elephants in India. Elephants in the Himalayan foothills from the Northwest to the Northeast of India are a single genetic cluster that diverged from other Indian populations more than 70,000 years ago. This makes the Northern (NW&NE) cluster the oldest and, hence, the most evolutionarily unique population.

Elephant to South of the Shencottah Gap need priority conservation attention to better understand census sizes, connectivity and genomic variation. Conservation and management action should focus on protecting remaining habitats, minimise unnatural deaths, examine the feasibility of restoring the link between NPG and SSG, and serious considerations about translocations into and out of this population. More genetic sampling of the SSG population as well as studies on inbreeding and possible associated phenotypes will be important to understand future fitness trajectories here.

Populations to the North and South of the Palghat Gap are distinct management units, and animals should be moved across this biogeographic divide only after very careful biological evaluations of the consequences. Further, given the high genetic load in these two management units, connectivity must be maintained within each of these two major landscapes across the gap.

The Ganges and Brahmaputra rivers in the north function as biogeographic divides, though incomplete, making elephants south of these rivers in Central India a unique management/conservation unit. Detailed and geographically widespread genetic studies in central India are important to better understand this management unit, in particular in the recent context of large-scale dispersals.

Overall, elephants across India face significant challenges because of a suite of factors, mainly negative human impacts, and we hope that conservation efforts can be bolstered by recognition of their unique evolutionary history. Using whole genome data (Weins et al. 2010, Toews and Brelsford 2012) as opposed to solely mitochondrial DNA (Vidya et al. 2009) can allow a better understanding of how Asian elephant evolution was shaped by Pleistocene climate cycles, and will respond to future climate change in the subcontinent (Kanagaraj et al. 2019). Previously researchers have suggested mixing lineages for conservation (provided risks of outbreeding depression are evaluated (Ralls et al. 2018), or assumptions can be made to ignore outbreeding depression (Powell 2023)). We find deep divergences between the populations of elephants. We recommend that maintaining historic connectivity between populations to allow natural dispersal between populations will be a better approach than translocation for conservation until risks of geneflow can be evaluated with empirical evidence.

While the impact of genetic variation on survival of small and isolated populations continues to be investigated, several recent studies highlight purging of deleterious allele load (Khan et al. 2021, Robinson et al. 2023 and the references therein). This study on Indian elephants reveals that while the number of deleterious alleles is low in small populations (such as SSG), the realized load is high because of higher homozygosity. Additionally, our data demonstrate that populations can have low (but unmasked) deleterious allele load due to their unique demographic histories.

Regardless, populations with homozygous deleterious mutation load should have high priority for conservation. The exact phenotypic and demographic trajectories of inbreeding depression and extinction will depend on the specific load, and maybe difficult to generalise across populations and species.

Our approach of using high-throughput genomic data allowed for unique and important conservation insights. Elephants in India present one of the first clear examples of serial founding of a large landscape at a subcontinental scale, with decreasing founder size from the source. Our data allows us to detect recent declines, potentially mediated by historic elephant captures on a large scale. Further, elephants provide an excellent system to better understand the interplay between genetic load and inbreeding in a set of five populations, a rare set up in endangered species. While on-ground conservation challenges for elephants remain mitigation of human-associated conflict and human infrastructure associated mortality, conservation genomics insights provide long-term conceptual guidance for future survival of these populations.

## Methods

### Sample collection

State Forest Department databases of captive elephants were examined from 6 states, to locate individual elephants that were captured or rescued from all of the known four major disjunct populations across India. Within these populations, individuals representing habitats across major barriers such as the Palghat and the Shencottah Gaps in Southern India, and the Ganges and the Brahmaputra rivers in Northern (Northwestern and Northeastern) India were also located. Blood samples and exact capture locations were collected from 28 such individuals and included in our final dataset (see Supplementary Table 1). Out of those samples, 12 were collected from the Southern eco-region, four were collected from the Central eco-region, three were collected from the Northwestern eco-region, and lastly nine were collected from the Northeastern eco-region. Out of the 12 samples collected from the Southern eco-region, six were collected from North of Palghat Gap, four from South of Palghat Gap (but North of Shencottah Gap), and two from South of Shencottah Gap. We additionally included genomic data from five more individuals obtained from online sources — one individual each from Borneo, Myanmar and Northeastern India, and three individuals from Southern India (two North of Palghat Gap and one South of Palghat Gap) (See Supplementary Table 1).

### DNA extraction, library preparation and sequencing

Genomic DNA was extracted from blood samples using Qiagen DNeasy Blood & Tissue Kit. The library preparation and whole genome resequencing was carried out at Medgenome INC. The DNA extraction and sequencing were carried out following Khan et al., 2020.

### Variant calling and filtering

The raw sequencing reads were trimmed using the default settings in TrimGalore- 0.4.5 (https://github.com/FelixKrueger/TrimGalore). The trimmed reads were mapped to the *Elephas maximus* reference genome (https://dnazoo.s3.wasabisys.com/index.html?prefix=Elephas_maximus/) using default settings of BWA *mem* (https://github.com/lh3/bwa). The mapped reads were converted to binary format and sorted using Samtools-1.9 (Li et al. 2009). The PCR and optical duplicates were marked using Picardtools MarkDuplicates (http://broadinstitute.github.io/picard) or Samtools-markdup. SNPs were identified using Strelka variant caller 2.9.10 (Saunders et al. 2012) for only the chromosomal scaffolds (scaffolds named HiC_scaffold_1 to HiC_scaffold_28). The variants were filtered using VcftoolsV13 (Danecek et al. 2011). We removed indels and retained only SNP loci with minimum phred scaled base quality 30, minimum genotype quality 30, a minimum minor allele count of 3, did not deviate from Hardy-Weinberg equilibrium with chi square p value of 0.05. All sites with the FILTER flag other than PASS were removed. Sites that showed mean depth across individuals more than the 97.5^th^ percentile and less than 2.5^th^ percentile or were missing in more than 20% individuals, were removed. We identified the sex chromosome scaffolds as

described in Armstrong et al. (2021) and removed the sex chromosome scaffold identified as HiC_scaffold_28.

### Population genetic structure

#### PCA

We first employed PCA to partition the data along their main axes of variation. The PCA was carried out using the *--pca* function in Plink (Purcell et al. 2007) based on the final filtered variants. The resulting eigenvectors were plotted in R using ggplot2.

#### ADMIXTURE

Thereafter, a maximum likelihood-based method, ADMIXTURE (Alexander et al. 2009) was employed to investigate the population genomic structure of Asian elephants. We estimated the number of clusters across *K* values ranging from 1 to 6, based on the number of clusters obtained from PCA. The evaluation of the most optimal number of clusters was carried out using EvalAdmix (GarcialJErill et al.

2020). EvalAdmix estimates the correlation of the residual matrices of the individuals from the Admixture analyses. The resulting correlation matrices were plotted in R.

#### qpgraph

We assigned a population to each individual based on the results from admixture. We converted the .vcf file to .ped format using VCFtools and then .ped to .bed format using Plink. Employed the *find_graphs* function in ADMIXTOOLS2 (https://github.com/uqrmaie1/admixtools) to automatically find the optimum graphs using various values for the number of admixture events. The model with zero admixture had the best statistical support as tested using the protocol suggested in ADMIXTOOLS2 with functions *qpgraph_resample_multi* and *compare_fits*.

### Demographic history

#### Recent Demographic History

We used the LD based GONE (Santiago et al. 2020) for estimating recent demography history up to a couple of hundred generations ago. We used the default settings of the parameters and set *PHASE* as 0. We plotted the results assuming a generation time of 31 years (Palkopoulou et al. 2018). This is a multiple sample analysis and requires several samples for accurate estimates. Hence, we performed this analysis only for the NW&NE and the NPG cluster which have at least nine samples each. The results from this analysis better predict recent events but are not very reliable for estimating events older than a hundred generations (Santiago et al. 2020).

#### Divergence time

Divergence time was estimated by contrasting various models using *fastsimcoal2.7*

(Excoffier et al. 2021) and hPSMC (Cahill et al. 2016).

We filtered the autosomal SNPs to remove sites that are in LD with r^2^ value of 0.5. This retained 80,684 SNPs. We estimated folded site frequency spectrum using easySFS (https://github.com/isaacovercast/easySFS) to subsample 9,4,6,4,2 diploid individuals from the NW&NE, CI, NPG, SPG and SSG populations using the options *--proj 18,8,12,8,4.* This allowed us to retain maximum number of non-missing SNPs for each of these populations while maximising the number of individuals retained. We tested scenarios presented in Figures 7a and 7b with constant population size and growth rate. We model two scenarios of divergence for both models in Figures 7a and 7b, one with advent of agriculture (7000 ya) as the maximum time of divergence and another with minimum divergence time around the last glacial maxima (17,500ya, Jones et al. 2017). We estimated parameters using fastsimcoal2.7 using options *-n 500,000 -m -M 0.001 -l 30 -L 60* as described in Armstrong et al. (2021). We then selected the best model using the best AIC values (Supplementary Table 8). We estimated confidence intervals using bootstrap. For this, we randomly sampled 80% of the loci from the dataset 10 times and estimated parameter with initial values declared using *-initvalues* and fastsimcoal2.7 options *-n 500,000 -m -M 0.001 -l 30 -L 60*.

**Figure7.**
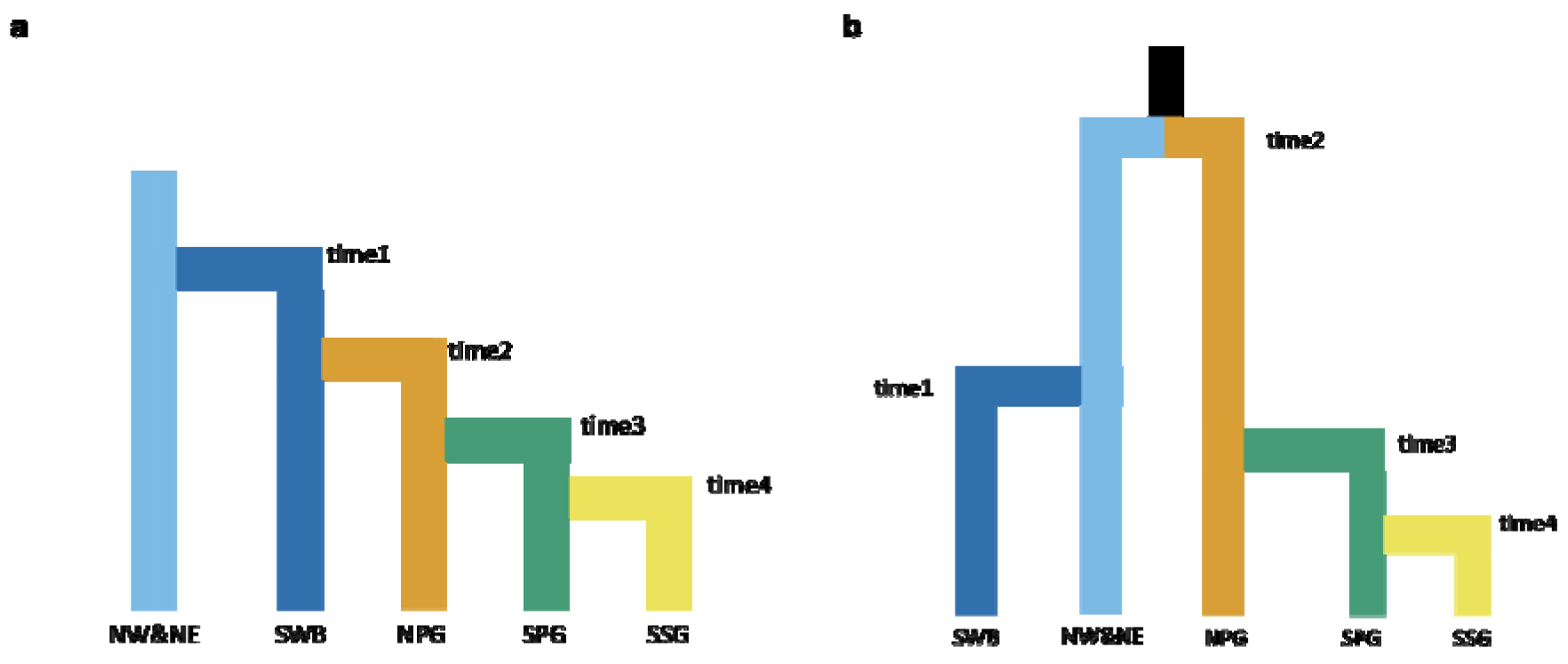
(a, b). Models tested on fastsimcoal2.7 using scenarios of recent divergence (less than 7,000 ya) and old divergence (more than 17,500 ya).

We also estimate divergence times between populations using hPSMC (Cahill et al. 2016). We used a single individual selected at random from each population (Supplementary Table 1). We created a pseudo-diploid individual using samples from two populations for each scaffold except for the sex chromosome scaffold

(https://github.com/jacahill/hPSMC). Then performed PSMC with the default settings. We plotted the results assuming a mutation rate of 5.3*10^−9/base/generation (Prado et al. 2023) and a generation time of 31 years. The time point where the effective population size estimated from the pseudo-diploid individual rises exponentially is the point where the two haplotypes do not coalesce and hence signify population divergence.

### Genetic diversity

#### Nucleotide diversity: pi

We randomly subsampled four individuals from NW&NE, CI, NPG and SPG sample sets and retained the SSG dataset (n=2) as is. We estimated the average number of pairwise differences per site for each population cluster as described in Wang et al. (2020). Briefly, we set the function *dosaf* to 1 ANGSD (Korneliussen et al. 2014) to estimate allele frequency likelihood for each site. The we used the used the function *realSFS* and estimated a folded site frequency spectrum. We set the function *doThetas* to 1 and estimated pi per site and used the *ThetaStat* function the summarise the pi value for each scaffold. We used only the chromosomal scaffolds from this summary and divided the “tP” values by the number of sites for each scaffold to obtain the average pi per site. Furthermore, we statistically compared the significance of the estimated values between populations (see Supplementary Table 4).

#### Heterozygous SNV encounter rate

We estimated the number of heterozygous SNPs for each individual using the *vcfstats* function of RTGtools (Cleary et al. 2015, https://github.com/RealTimeGenomics/rtg-tools). We then divided the number of heterozygous sites by the total number of sites genotyped for the individual and multiplied by 10^6 to obtain SNV encounter rate/Mb.

### Inbreeding

#### FROH

We estimated runs of homozygosity (ROH) using the *roh* function in BCFtools version 1.3 (Narasimhan et al. 2016) as described previously (Shukla et al. 2023). We classified the ROH stretches into three size classes: the more than 0.1 Mb (100 Kb) stretches show cumulative inbreeding due to old and recent bottlenecks, the more than 1Mb stretches show cumulative inbreeding in the recent past and the more than 10Mb stretches show recent inbreeding. FROH was estimated as described previously (Khan et al. 2021).

#### Shared IBD stretches

We used IBDseq (Browning & Browning, 2013) to detect the stretches of genome that are identical-by-descent in pairs of individuals. For each pair of individuals in a population, we summed the lengths of IBD stretches longer than 0.1Mb and 10Mb and divided by the total autosomal length.

### Genetic load

#### Filtering variant sites

We filtered the raw variants obtained from Strelka variant caller 2.9.10 again in VCFtools using the parameters with minimum phred scaled base quality 30, minimum genotype quality 30, minor allele count 1, FILTER flag as PASS. We also removed sited with -0.5>Fis>0.95 as described previously (Khan et al. 2021) and removed sites missing in more than 20% of the individuals. We filtered out the sex chromosomes as well. We call this set of loci as deleterious_allele_set.

#### Identifying ancestral alleles

We defined the ancestral allele as the allele present most commonly in sister species of Asiatic elephants. For this we used the genome of African elephant (genbank accession: GCA_000001905.1) and dugong (genbank accession: GCA_015147995). We converted these assemblies to 100bp long FASTQ reads and mapped to the Asiatic elephant reference genome and removed reads mapping to multiple regions as described in Khan et al. (2021). We converted the mapped reads to consensus fasta files and estimated the majority allele at a locus. We further filtered these alleles with the genome of Bornean elephant (SRA accession: ERR2260499). Any

loci where the Bornean elephant was not homozygous for the ancestral allele was removed from the analysis thus ensuring that the derived alleles are new to the population which is an important assumption for the identification of deleterious alleles (Khan 2023).

#### Derived indel load

We further filtered the loci in the deleterious_allele_set using the *--keep-only-indels* tag in VCFtools. We annotated this set of loci with the Asiatic elephant genome annotation (https://dnazoo.s3.wasabisys.com/index.html?prefix=Elephas_maximus/), consisting of 26246 coding regions, using Ensembl Variant Effect Predictor (VEP, McLaren et al. 2016). For indels, the ancestral state cannot be determined using distant species. Hence, the indel allele that was homozygous in the Bornean elephant was referred to as the ancestral allele and any indel site where the Bornean elephant was not homozygous, was removed from the analysis. We then estimated the number of indels predicted to cause transcript_ablation, splice_donor_variant, splice_acceptor_variant, stop_gained, frameshift_variant, inframe_insertion, inframe_deletion, splice_region_variant were classified as LoF causing indels. We also counted the derived indels in the intergenic regions of the genome. We divided the number of LoF indels with the number of intergenic indels to control for differences in depth leading to missingness in the data.

#### Derived LoF load

We filtered the loci in the deleterious_allele_set to remove indels and that had mean depth across individuals more than the 97.5^th^ percentile and less than 2.5^th^ percentile. This set was annotated on VEP and LoF mutations were identified as described earlier. We then counted the number of derived LoF alleles and divided them with the number of synonymous mutations.

#### Derived missense load

We used the same set of loci used for estimating the LoF load but chose those set of loci that caused non-synonymous changes in the genome and followed the same procedure as described for the LoF load.

## Acknowledgements

This work was supported by a Department of Biotechnology grant [BT/PR23223/BCE/8/1490/2018] awarded to RS and UR. Permissions for sample collection were granted in letters C-66011/02/2019 dated 09/07/2019 (West Bengal), 1729/5-6 dated 09/12/2020 (Uttarakhand), 4838/2019-CWW/WL10 dated 06/10/2019 (Kerala), PCCF(WL)/E2/CR-79/2018-19 dated 26/05/2018 (Karnataka), WL/FG31/Technical Committee/2019 dated 26/08/2019 (Assam) and CWL/G/173/2018-19/Pt.VII/1695-702 dated 10/10/2019 (Arunachal Pradesh). We would like to thank the respective forest veterinary doctors of all the states, particularly Dr. Aditi Sharma, Dr. Arun Zachariah and Dr. Shweta Mandal. RS was supported as National Science Chair by Science and Engineering Research Board (SERB), Government of India. KMG is supported from the DBT-Ramalingaswami Fellowship (No. BT/HRD/35/02/2006). Gratitude to NCBS IT team support during data analysis; NCBS data cluster used is supported under project no. 12-R&D-TFR- 5.04-0900, Department of Atomic Energy, Government of India). VV Robin and Ansil BR provided helpful comments on drafts of the manuscript. Gratitude to field assistants for their continuous presence in the field and assisting in sample collection for this work.

## Data availability

All sequencing data have been deposited in BioProject SUB13810487 in NCBI.

