## Supplementary materials for "Serial colonization shapes genetic variation and defines conservation units in Asian elephants"

**Supplementary Methods:**

**LD filtering:**

We estimated the loci in LD using PLINK (Purcell et al. 2007) *–indep-pairwise* in 50kb windows with a 10 SNP step size for an r2 threshold of 0.5. Using the out produced from PLINK, the set of SNPs were filtered using vcftools (Danecek et al. 2011).

**EvalAdmix:**

We evaluated the ADMIXTURE plots by quantifying the residual genotypes observed in the data and predicted from the model suggested by various K values using evalAdmix (<http://www.popgen.dk/software/index.php/EvalAdmix>, Garcia-Eril et al. 2020). We used evaladmix as explained in the user manual and then plotted the results. The K value with the least cross-correlation between residuals was chosen.

**Neighbour Joining Tree:**

A Neighbour Joining (NJ) Tree was constructed LD pruned dataset. We converted the VCF file to phylip file using vcftophylip.py (https://github.com/edgardomortiz/vcf2phylip). The tree was constructed in PAUP4.0a (Swofford, 2003).

**FST:**

We estimated pairwise FST between populations using VCFtools. Both mean and weighted FST were estimated in order to capture the relative separation of the populations.

**F_3_:**

We estimated F_3_ statistics to test for geneflow between various pairs of population under various three population model of trees. ADMIXTOOLS2 was used to estimate F_3_ values for all possible three population models in our data. The function *qp3pop* and *f3* were used.

**Statistical estimate of Nucleotide diversity (pi) & heterozygous SNV encounter rate per Mb:**

The pi values were statistically compared between each population. We carried out Shapiro-Wilk normality test and Bartlett test of homogeneity of variances to determine if ANOVA can be carried out on our dataset. Then we carried out a one-way ANOVA test implemented in R to estimate if the median values of these populations are significantly different. Thereafter tukey's hsd was conducted to identify the groups that were significantly different. Similarly, we carried out a GLM with poissson distribution to test whether the difference in heterozygous SNV encounter rate per Mb was different between the populations. All the analyses has been carried out in R.

**Shared Alleles:**

We randomly chose two individuals from each population. We used SnpCountCU (<https://github.com/JingfangSI/SnpCountCU>) to count the number of SNPs common among populations with default settings. We then estimated percent SNPs shared across populations genome-wide.

**Old demographic History**

We used the coalescent based SMC++ (Terhorst et al. 2017) method for estimating older demographic history. We used the protocol described for the implementation of the method (<https://github.com/popgenmethods/smcpp>) as is. We set the mutation rate to 5.3*10^-9/base/generation (Prado et al. 2023) and a generation time of 31 years. The estimates from this method are reliable for estimating demographic events between a hundred generations ago to forty thousand generations ago.

**Gene Enrichment:**

The symbols of genes containing missense and LOF mutations were output from VEP. The genes with unknown functions were removed from the lists. The genes with missense mutations and the total genes in the elephant genome were uploaded to Webgestalt (<https://www.webgestalt.org/>). Humans were chosen as the organism of interest, Over-Representation Analysis was chosen as the method of interest. The geneontology database with biological process, cellular component and molecular function was chosen as the database. We set minimum 3 genes per category and the run the analysis at default for other settings.

**Mitogenome analyses:** TrimGalore-0.4.5 was employed to trim the raw reads, which were then mapped to the *Elephas maximus* whole mitogenome (https://www.ncbi.nlm.nih.gov/nucleotide/NC_005129.2), using BWA *mem*. We converted the mapped reads to binary format and sorted them using Samtools-1.9. Thereafter we used the Picardtools MarkDuplicate function or Samtools-markdup to mark the PCR duplicates. The bam files were converted to consensus fasta files using ANGSD. The fasta files were aligned using Clustal Omega (https://www.ebi.ac.uk/Tools/msa/clustalo/). A TCS haplotype network (Clement et al., 2002) was constructed based on the alignment, implemented in popart-1.7 (Leigh & Bryant, 2015).

Supplementary Table 1: Sample information of the Asian elephants (F = female, M = male) sampled along with location of capture/rescue and sequencing depth

| Name of the elephant | Serial No. | Population | Capture/rescue Location | Source | Depth of sequencing |
| --- | --- | --- | --- | --- | --- |
| Varalakshmi (F) | KA_MTG_02 | NPG | Hebbala – Kodagu, Karnataka | Current Study | 18.93 |
| Sri Ranga (M) | KA_MTG_03 | NPG | Ramnagara – Bangalore rural, Karnataka | Current Study | 15.84 |
| Bheema (M) | KA_MTG_08 | NPG | Bheemankatte – Nagarhole, Karnataka | Current Study | 14.79 |
| Ramaiah (M) | KA_MTG_05 | NPG | Umbilibetta – Hassan, Karnataka | Current Study | 11.84 |
| Dhruva (M) | KA_MTG_11 | NPG | Hassan, Karnataka | Current Study | 19.57 |
| Jayaprakash (M) | NA | NPG | Bandipur, Karnataka | Reddy et al. 2015 | 14.80 |
| Krishna (M) | KA_RM_01 | NPG | Bandipur camp, born to Chaitra (captive female) and a sired by a wild bull | Current Study | 21.43 |
| Bheem (M) | KA_MTG_09 | NPG | Savanthavadi, Maharashtra | Current Study | 11.95 |
| Parvathy (F) | KL_KD_04 | NPG | Kozhikode, Kerala | Current Study | 28.23 |
| Anjana (F) | KL_KD_03 | SPG | Kodanadu Range, Malayattur Division, Kerala | Current Study | 12.49 |
| Asha (F) | NA | SPG | Kerala | Palkopoulou et al. (2018) | 31.93 |
| Jayashree (F) | KL_KKT_01 | SPG | Kodanadu Range, Malayattur Division, Kerala | Current Study | 19.91 |
| Minna (F) | KL_KKT_05 | SPG | Puyankutti, Kothamangalam Division, Kerala | Current Study | 22.80 |
| Podichi (F) | KL_KKT_04 | SSG | Podiyakala, Peppara Range, Kerala | Current Study | 20.73 |
| Raja (M) | KL_KKT_06 | SSG | Kottur, Kerala | Current Study | 20.12 |
| Kaberi (F) | WB_GMD_10 | Central | Garseta Range, Midnapore, West Bengal | Current Study | 25.42 |
| Aranya (M) | WB_GMG_06 | Central | Burdwan Division, West Bengal | Current Study | 26.57 |
| Rami (F) | WB_GMM_01 | Central | West Midnapore, West Bengal | Current Study | 12.80 |
| Balaram (M) | WB_JPK_03 | Central | Bagti Mata Range, Purulia Division, West Bengal | Current Study | 19.53 |
| Bolanath (M) | WB_GMG_03 | Central | Kaliaganj, North Dinajpur, West Bengal | Current Study | 20.83 |
| Balasundar (M) | WB_JPN_02 | Northeast | Balason River, Bagdogra Range, West Bengal | Current Study | 10.58 |
| Tamuk (M) | AP_PKT_02 | Northeast | Namsai Forest Division, Arunachal Pradesh | Current Study | 16.57 |
| Babu (M) | AP_PKS_02 | Northeast | Pakke Tiger Reserve, Arunachal Pradesh | Current Study | 19.97 |
| Lokmuti (F) | AP_PKS_03 | Northeast | Pasighat Division, Arunachal Pradesh | Current Study | 18.41 |
| Maya | NA | Northeast | Assam | Palkopoulou et al. (2018) | 29.31 |
| Samrat (MK) | AS_KZC_02 | Northeast | Dala Mara Reserve Forest, Assam | Current Study | 16.85 |
| Bishnu (MK) | AS_KZC_09 | Northeast | Nagaon Division, Assam | Current Study | 20.25 |
| Konwari (F) | AS_KZE_01 | Northeast | Kaziranga National Park, Assam | Current Study | 19.43 |
| Phaguni (F) | AS_KZE_02 | Northeast | Sonitpur East Division, Assam | Current Study | 18.86 |
| Gomti (F) | UK_JC_02 | Northwest | Bahraich, Uttar Pradesh | Current Study | 18.67 |
| Pavanpari (F) | UK_JC_03 | Northwest | Bahraich, Uttar Pradesh | Current Study | 17.67 |
| Raja (M) | UK_RJI_01 | Northwest | Rajaji/Haridwar, Uttarakhand | Current Study | 17.11 |
| Moola | NA | Myanmar |  | Palkopoulou et al. (2018) | 30.58 |
| Chendra | NA | Borneo |  | Palkopoulou et al. (2018) | 32.2 |

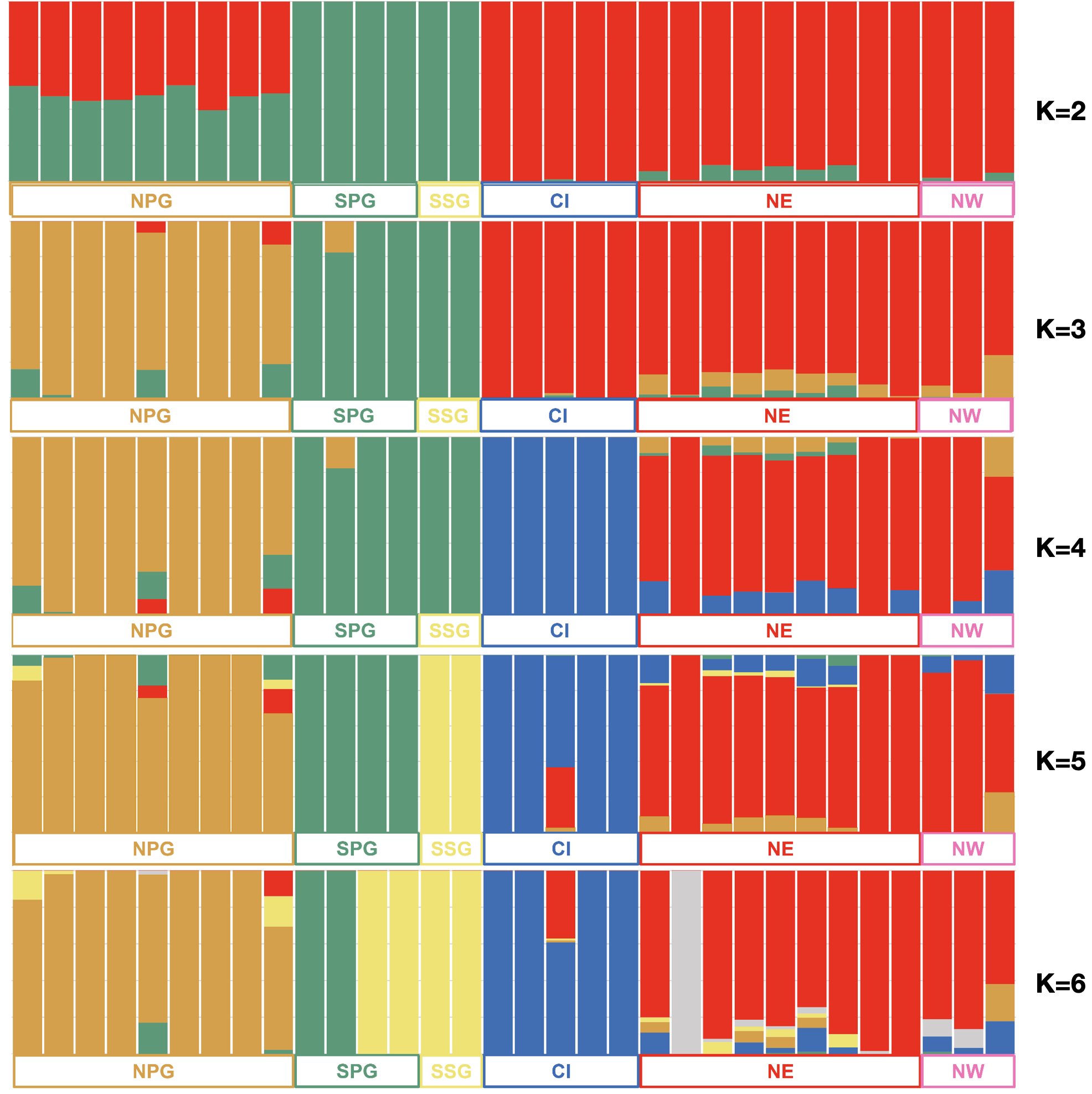

Supplementary Figure 1: ADMIXTURE plots from K=2 to K=6

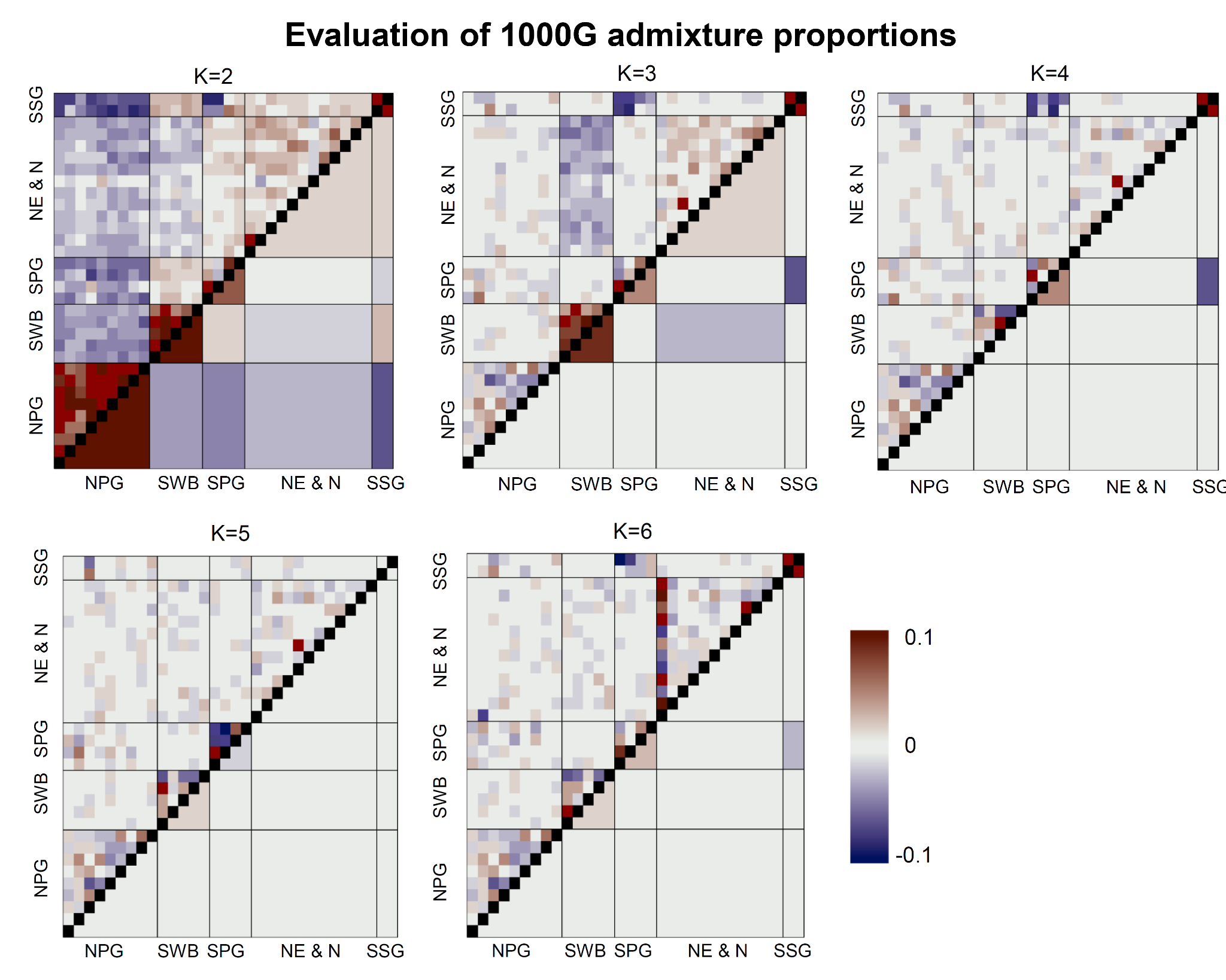

Supplementary Figure 2: Evaluation of residuals from structure plots. Evaladmix K=2 to K=6

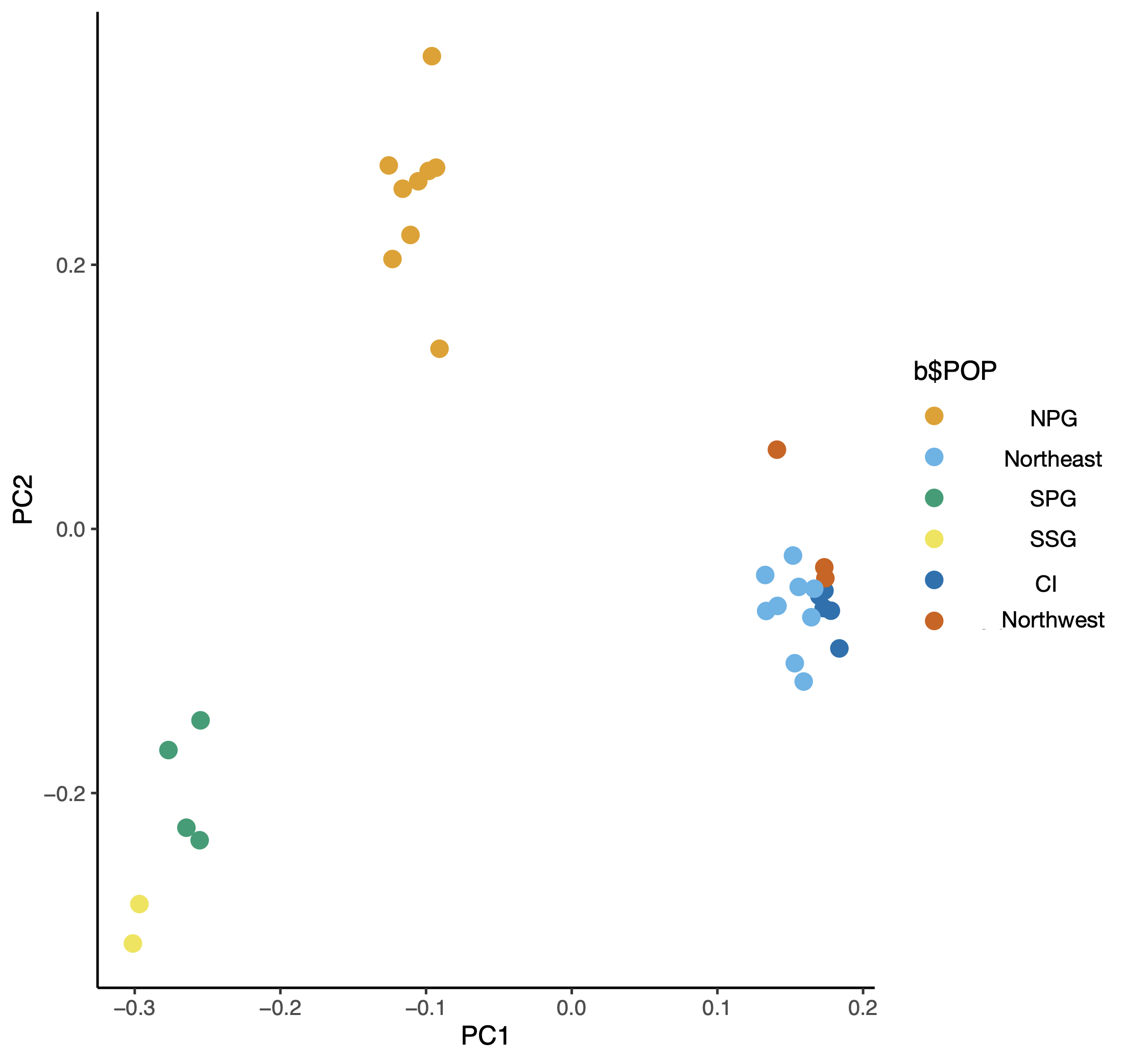

LEGEND

Supplementary Figure 3: PCA plot of all individuals after removing sites in LD

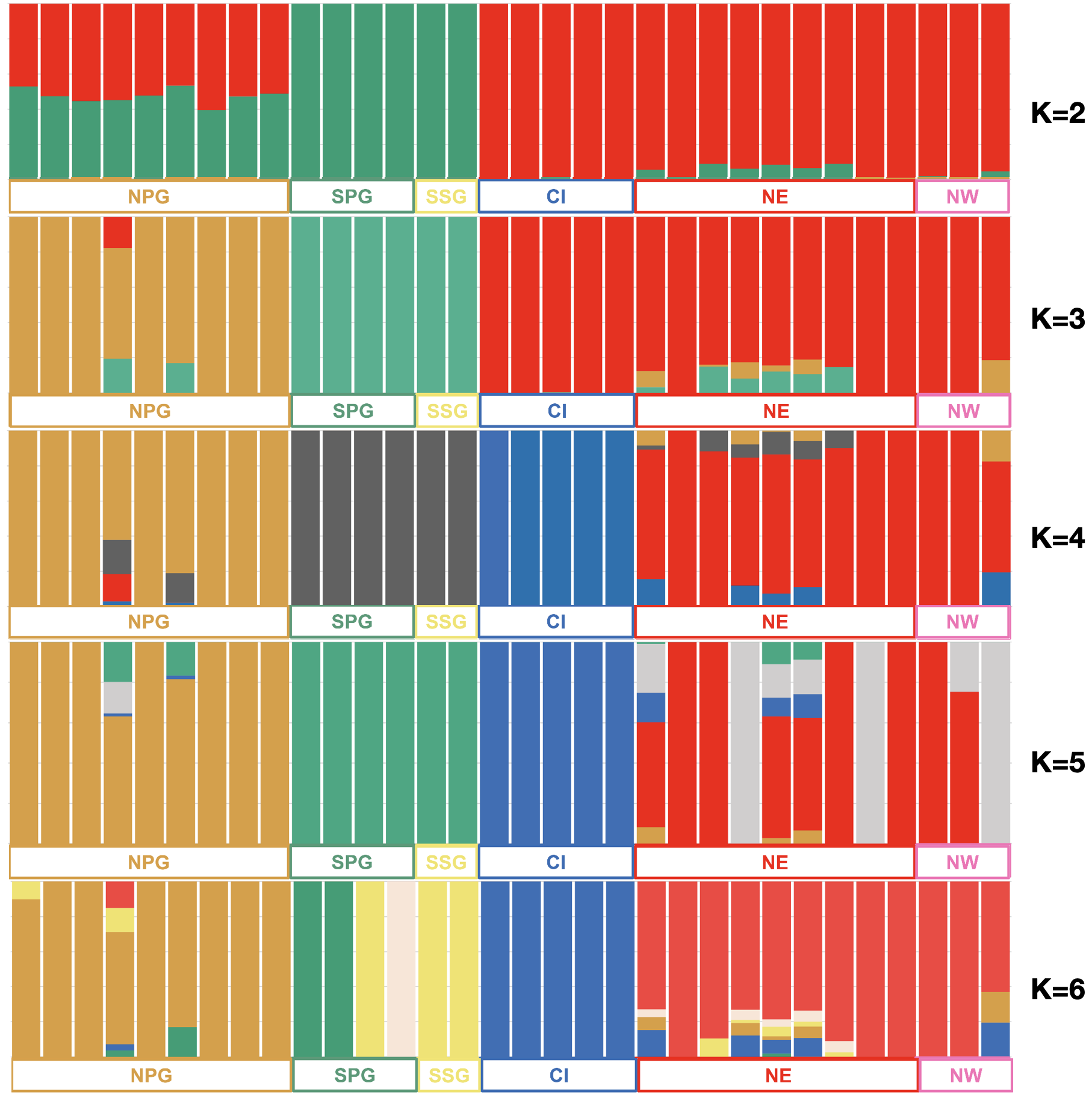

Supplementary Figure 4: ADMIXTURE plots using LD filtered SNPs from K=2 to K=6

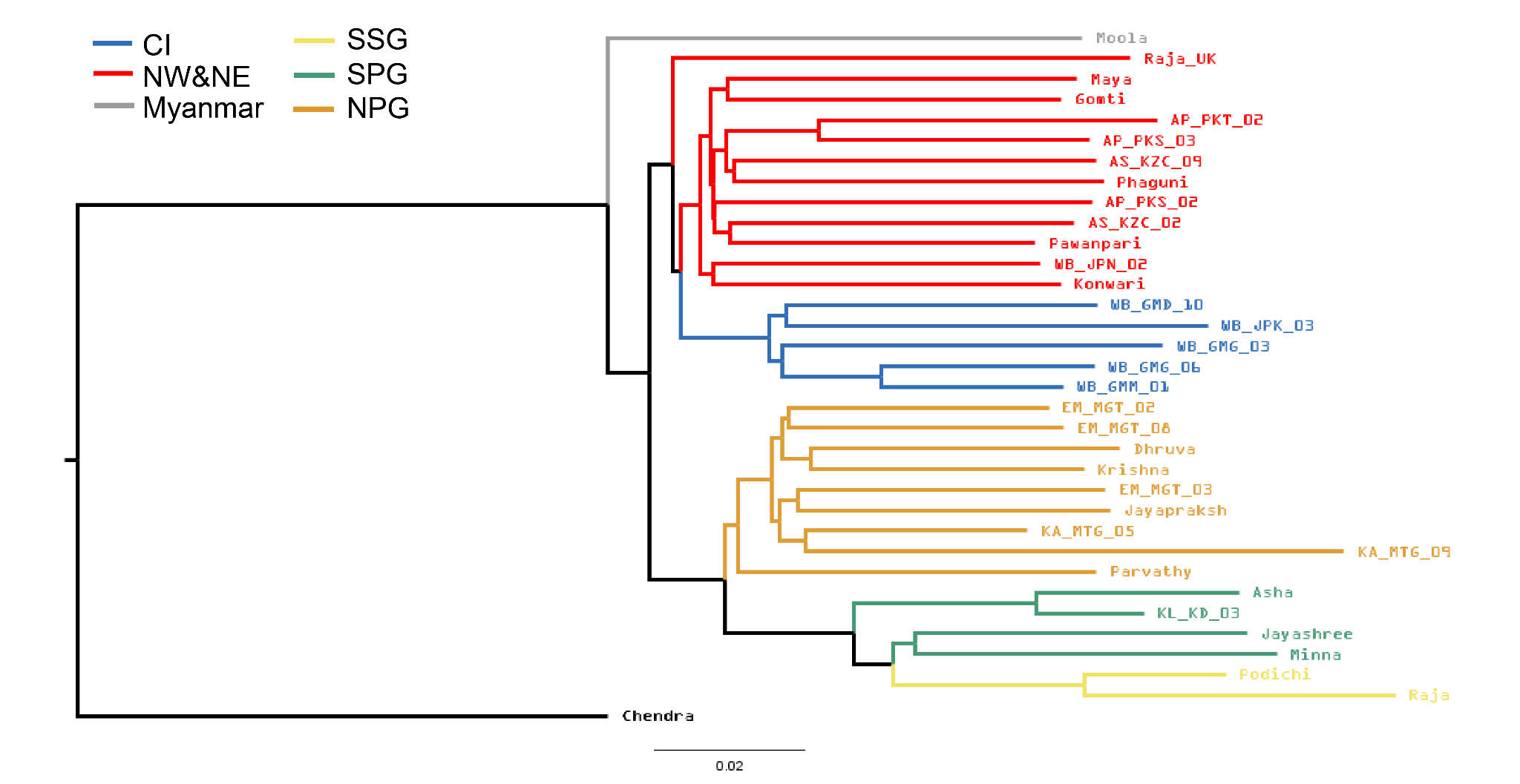

Supplementary Figure 5: A neighbour joining tree constructed from the LD pruned SNPs

Supplementary Figure 6: raw qpgraph with K 0

Supplementary Table 2: FST values between populations

| **Population 1** | **Population 2** | **Mean FST** | **Weighted FST** |
| --- | --- | --- | --- |
| SSG | SPG | 0.017 | 0.071 |
| SSG | NPG | 0.054 | 0.126 |
| SSG | CI | 0.100 | 0.184 |
| SSG | NW & NE | 0.070 | 0.135 |
| SPG | NPG | 0.058 | 0.091 |
| SPG | CI | 0.104 | 0.153 |
| SPG | NW & NE | 0.078 | 0.112 |
| NPG | CI | 0.066 | 0.090 |
| NPG | NW & NE | 0.048 | 0.060 |
| CI | NW & NE | 0.018 | 0.031 |

Supplementary Table 3: F3 statistics to evaluate geneflow between populations 1 and 2 under various three population trees. Negative values indicate recent geneflow and p values indicate significance

| **pop1** | **pop2** | **pop3** | **F_3_** | **se** | **z** | **p** |
| --- | --- | --- | --- | --- | --- | --- |
| Burma | NW&NE | Borneo | -0.00188 | 0.00185 | -1.01530 | 0.30997 |
| N&NE | CI | Burma | -0.00033 | 0.00045 | -0.73076 | 0.46492 |
| Burma | CI | Borneo | 0.00019 | 0.00180 | 0.10880 | 0.91336 |
| N&NE | NPG | CI | 0.00020 | 0.00033 | 0.60989 | 0.54193 |
| N&NE | SPG | CI | 0.00063 | 0.00042 | 1.48446 | 0.13769 |
| N&NE | SSG | CI | 0.00098 | 0.00050 | 1.94872 | 0.05133 |
| SPG | SSG | NW&NE | 0.00099 | 0.00070 | 1.35343 | 0.17592 |
| Burma | NPG | Borneo | 0.00115 | 0.00184 | 0.62649 | 0.53099 |
| SPG | SSG | NPG | 0.00126 | 0.00074 | 1.70940 | 0.08738 |
| SPG | SSG | CI | 0.00134 | 0.00078 | 1.72877 | 0.08385 |
| SPG | SSG | Burma | 0.00154 | 0.00097 | 1.58915 | 0.11202 |
| N&NE | CI | Borneo | 0.00174 | 0.00055 | 3.19063 | 0.00142 |
| Burma | NW&NE | SSG | 0.00188 | 0.00154 | 1.21856 | 0.22301 |
| Burma | NW&NE | SPG | 0.00244 | 0.00154 | 1.57440 | 0.11539 |
| NPG | SSG | NW&NE | 0.00289 | 0.00053 | 5.42125 | 5.92E-08 |
| SPG | SSG | Borneo | 0.00301 | 0.00089 | 3.38730 | 0.00070 |
| NPG | SPG | NW&NE | 0.00316 | 0.00042 | 7.52787 | 5.16E-14 |
| Burma | SSG | CI | 0.00319 | 0.00153 | 2.08773 | 0.03682 |
| Burma | SPG | CI | 0.00339 | 0.00158 | 2.14759 | 0.03175 |
| NPG | SPG | CI | 0.00358 | 0.00054 | 6.60688 | 3.93E-11 |
| NPG | SSG | CI | 0.00366 | 0.00070 | 5.24002 | 1.61E-07 |
| Burma | SPG | Borneo | 0.00422 | 0.00197 | 2.14197 | 0.03219 |
| N&NE | NPG | Burma | 0.00446 | 0.00051 | 8.69273 | 3.54E-18 |
| Burma | NW&NE | NPG | 0.00530 | 0.00149 | 3.56350 | 0.00036 |
| Burma | SSG | Borneo | 0.00569 | 0.00185 | 3.06696 | 0.00216 |
| Burma | NPG | CI | 0.00583 | 0.00145 | 4.01520 | 5.94E-05 |
| NPG | SPG | Burma | 0.00602 | 0.0006 | 10.0609 | 8.23E-24 |
| NPG | SSG | Burma | 0.00630 | 0.00075 | 8.40880 | 4.14E-17 |
| N&NE | SPG | Burma | 0.00732 | 0.00066 | 11.09370 | 1.35E-28 |
| N&NE | NPG | Borneo | 0.00749 | 0.00057 | 13.01029 | 1.07E-38 |
| N&NE | SSG | Burma | 0.00787 | 0.00088 | 8.95084 | 3.53E-19 |
| CI | Borneo | NW&NE | 0.00887 | 0.00062 | 14.26087 | 3.84E-46 |
| NPG | SPG | Borneo | 0.00909 | 0.00067 | 13.63169 | 2.60E-42 |
| CI | NW&NE | SSG | 0.00965 | 0.00058 | 16.53634 | 2.01E-61 |
| CI | NW&NE | SPG | 0.01000 | 0.00050 | 19.97498 | 9.09E-89 |
| Burma | NW&NE | CI | 0.01008 | 0.00147 | 6.87694 | 6.11E-12 |
| CI | NW&NE | NPG | 0.01042 | 0.00049 | 21.20050 | 9.44E-100 |
| NPG | SSG | Borneo | 0.01084 | 0.00083 | 13.00590 | 1.13E-38 |
| CI | Burma | NW&NE | 0.01095 | 0.00056 | 19.27223 | 9.19E-83 |
| NW&NE | Borneo | Burma | 0.01163 | 0.00104 | 11.21090 | 3.60E-29 |
| NW&NE | SPG | Borneo | 0.01342 | 0.00072 | 18.58260 | 4.44E-77 |
| NPG | Borneo | NW&NE | 0.01423 | 0.00058 | 24.55398 | 3.92E-133 |
| CI | Burma | NPG | 0.01520 | 0.00079 | 19.20251 | 3.53E-82 |
| NW&NE | SSG | Borneo | 0.01544 | 0.00088 | 17.56060 | 4.94E-69 |
| NPG | CI | Borneo | 0.01577 | 0.00066 | 23.2323 | 2.15E-119 |
| CI | Borneo | NPG | 0.01616 | 0.00082 | 19.77021 | 5.37E-87 |
| Burma | NPG | SSG | 0.01625 | 0.00167 | 9.72716 | 2.31E-22 |
| Burma | NPG | SPG | 0.01654 | 0.00172 | 9.63689 | 5.59E-22 |
| NPG | CI | Burma | 0.01673 | 0.00071 | 23.52826 | 2.10E-122 |
| NPG | Burma | NW&NE | 0.01726 | 0.00053 | 32.34937 | 1.42E-229 |
| CI | Burma | SPG | 0.01764 | 0.00084 | 20.95863 | 1.57E-97 |
| CI | Burma | SSG | 0.01784 | 0.001137 | 15.68768 | 1.84E-55 |
| NW&NE | NPG | SPG | 0.01856 | 0.00042 | 44.59306 | 0 |
| NW&NE | NPG | SSG | 0.01883 | 0.00050 | 37.62919 | 0 |
| SSG | Borneo | SPG | 0.02053 | 0.00132 | 15.51302 | 2.83E-54 |
| CI | Borneo | Burma | 0.02084 | 0.00111 | 18.75989 | 1.61E-78 |
| NPG | Borneo | Burma | 0.02140 | 0.00104 | 20.55563 | 6.85E-94 |
| NPG | CI | NW&NE | 0.02151 | 0.00045 | 47.26812 | 0 |
| CI | Borneo | SPG | 0.02167 | 0.00087 | 24.94852 | 2.22E-137 |
| SSG | Burma | SPG | 0.02200 | 0.00138 | 15.95591 | 2.59E-57 |
| SSG | CI | SPG | 0.02220 | 0.00136 | 16.29903 | 1.00E-59 |
| SSG | NPG | SPG | 0.02228 | 0.00128 | 17.43500 | 4.48E-68 |
| SSG | NW&NE | SPG | 0.02255 | 0.00126 | 17.95148 | 4.67E-72 |
| SPG | Borneo | NPG | 0.02272 | 0.00081 | 28.04610 | 4.46E-173 |
| CI | Borneo | SSG | 0.02334 | 0.00105 | 22.13605 | 1.42E-108 |
| SPG | Burma | NPG | 0.02579 | 0.00075 | 34.23444 | 7.43E-257 |
| SPG | CI | NPG | 0.02822 | 0.00068 | 41.44365 | 0 |
| CI | NPG | SSG | 0.02827 | 0.00081 | 34.84090 | 5.85E-266 |
| CI | NPG | SPG | 0.02835 | 0.00061 | 46.62962 | 0 |
| SPG | NW&NE | NPG | 0.02865 | 0.00068 | 42.17110 | 0 |
| NPG | SPG | SSG | 0.03055 | 0.00085 | 36.01578 | 4.74E-284 |
| SPG | Borneo | NW&NE | 0.03379 | 0.00089 | 37.99739 | 0 |
| SPG | CI | Borneo | 0.03491 | 0.00089 | 39.08213 | 0 |
| SPG | Borneo | Burma | 0.03810 | 0.00122 | 31.16675 | 3.01E-213 |
| SPG | CI | Burma | 0.03893 | 0.00092 | 42.35525 | 0 |
| SPG | Burma | NW&NE | 0.03989 | 0.00086 | 46.32849 | 0 |
| Burma | SPG | SSG | 0.04078 | 0.00178 | 22.84087 | 1.80E-115 |
| SSG | Borneo | NPG | 0.04199 | 0.00150 | 27.90329 | 2.43E-171 |
| NW&NE | SPG | SSG | 0.04621 | 0.00090 | 51.27922 | 0 |
| SSG | Burma | NPG | 0.04653 | 0.00147 | 31.73058 | 5.88E-221 |
| SPG | CI | NW&NE | 0.04658 | 0.00079 | 58.91368 | 0 |
| SSG | CI | NPG | 0.04916 | 0.00152 | 32.40037 | 2.71E-230 |
| SSG | NW&NE | NPG | 0.04994 | 0.00142 | 35.02841 | 8.31E-269 |
| SSG | Borneo | NW&NE | 0.05333 | 0.00169 | 31.62642 | 1.60E-219 |
| SSG | CI | Borneo | 0.05410 | 0.00178 | 30.34229 | 3.18E-202 |
| CI | SPG | SSG | 0.05523 | 0.00099 | 55.50352 | 0 |
| SSG | Borneo | Burma | 0.05709 | 0.00200 | 28.47868 | 2.15E-178 |
| SSG | CI | Burma | 0.05959 | 0.00174 | 34.15433 | 1.15E-255 |
| SSG | Burma | NW&NE | 0.06090 | 0.00166 | 36.64674 | 5.16E-294 |
| SSG | CI | NW&NE | 0.06779 | 0.00163 | 41.66813 | 0 |
| Borneo | Burma | SSG | 0.16071 | 0.00205 | 78.27633 | 0 |
| Borneo | Burma | SPG | 0.16218 | 0.00184 | 87.98263 | 0 |
| Borneo | SSG | CI | 0.16371 | 0.00181 | 90.61025 | 0 |
| Borneo | NW&NE | SSG | 0.16447 | 0.00168 | 97.76815 | 0 |
| Borneo | Burma | NPG | 0.16525 | 0.00174 | 94.72844 | 0 |
| Borneo | SPG | CI | 0.16538 | 0.00162 | 101.90517 | 0 |
| Borneo | Burma | CI | 0.16621 | 0.00174 | 95.27640 | 0 |
| Borneo | NW&NE | SPG | 0.16650 | 0.00155 | 107.08806 | 0 |
| Borneo | Burma | NW&NE | 0.16828 | 0.00161 | 104.33706 | 0 |
| Borneo | NPG | CI | 0.17088 | 0.00150 | 114.11685 | 0 |
| Borneo | NW&NE | NPG | 0.17243 | 0.00144 | 119.36752 | 0 |
| Borneo | NPG | SSG | 0.17581 | 0.00183 | 96.27270 | 0 |
| Borneo | NPG | SPG | 0.17756 | 0.00170 | 104.51561 | 0 |
| Borneo | NW&NE | CI | 0.17817 | 0.00140 | 127.35586 | 0 |
| Borneo | SPG | SSG | 0.19727 | 0.00201 | 98.10156 | 0 |

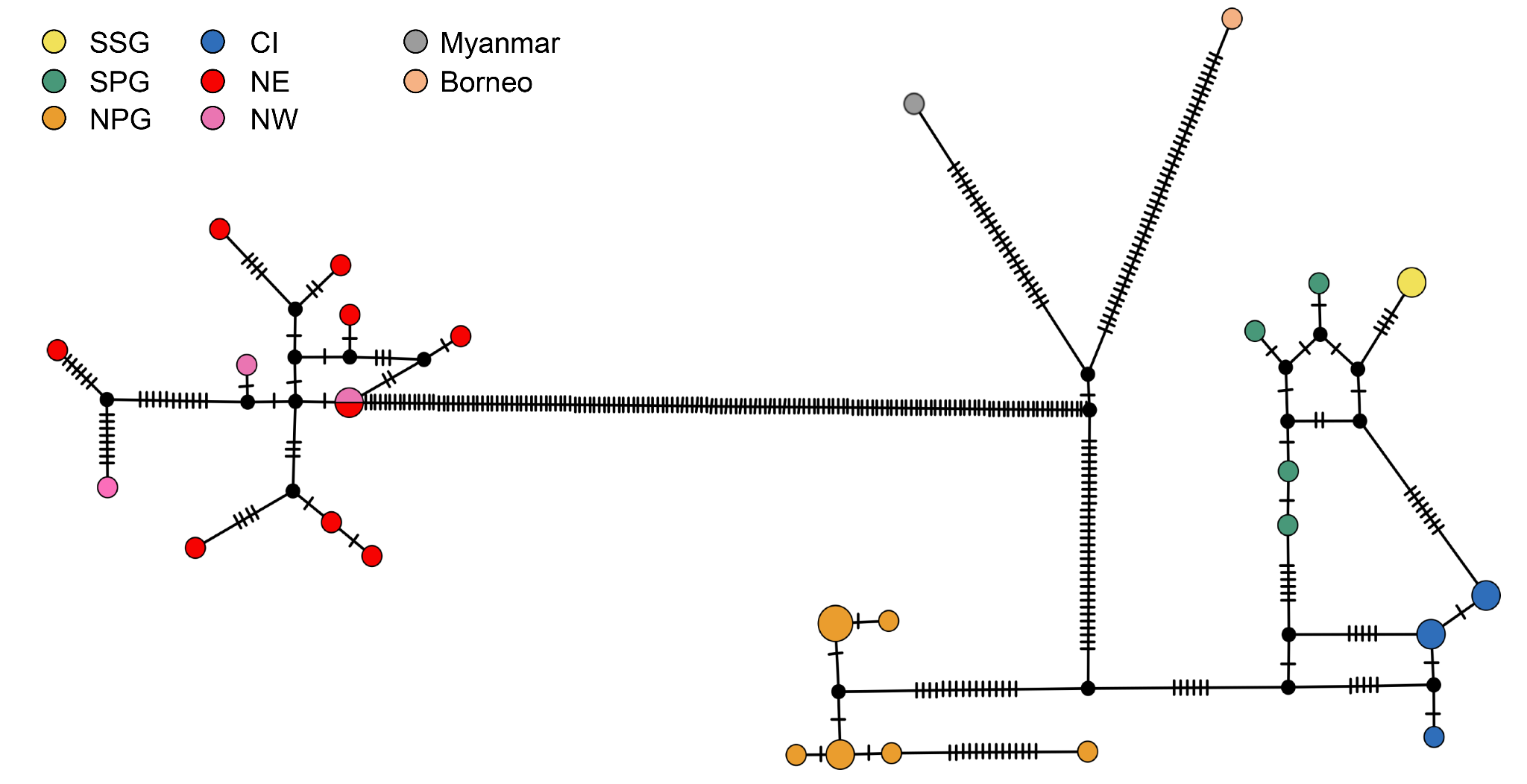

Supplementary Figure 7: Mitogenome network of Asiatic elephants; The size of the circles represent the number of individuals constituting that particular haplotype; the dashes represent mutational steps; the length of the lines represent one haplotype to another (not to scale).

Supplementary Table 4: Statistical estimation of Pi (Tukey multiple comparisons of means)

|  | NW & NE | CI | NPG | SPG | SSG |
| --- | --- | --- | --- | --- | --- |
| NW & NE | # | 4.19807e-05 | -1.20715e-04 | -1.37241e-04 | -1.06298e-04 |
| CI | 0.62898 | # | 1.62696e-04 | 1.79222e-04 | 1.48278e-04 |
| NPG | 0.00090*** | 2.7e-06*** | # | -1.65263e-05 | 1.44170e-05 |
| SPG | 0.00010*** | 2.0e-07*** | 0.98164 | # | 3.09433e-05 |
| SSG | 0.00490** | 2.27e-05*** | 0.98900 | 0.84025 | # |

Shapiro-Wilk normality test: W = 0.98258, p-value = 0.08228

Bartlett test of homogeneity of variances: Bartlett's K-squared = 1.34, df = 4, p-value = 0.8546

One-way ANOVA: F value = 14.28, p-value = 1.09e-09

Signif. codes: 0 ‘***’ 0.001 ‘**’ 0.01 ‘*’ 0.05 ‘.’

Supplementary Table 5: Statistical estimation of heterozygous SNV per Mb

|  | NW & NE | CI | NPG | SPG | SSG |
| --- | --- | --- | --- | --- | --- |
| NW & NE |  | -0.13149 | -0.13854 | -0.25527 | -0.39637 |
| CI | <2e-16 *** |  | -0.00705 | -0.12378 | -0.26488 |
| NPG | <2e-16 *** | 0.81979 |  | -0.11673 | -0.25782 |
| SPG | 2e-16 *** | 2e-16 *** | 2e-16 *** |  | -0.14110 |
| SSG | 2e-16 *** | 2e-16 *** | 2e-16 *** | 2e-16 *** |  |

Signif. codes: 0 ‘***’ 0.001 ‘**’ 0.01 ‘*’ 0.05 ‘.’ 0.1 ‘ ’ 1

Supplementary Table 6: Number of shared alleles

|  |  | Unique | | | | |  |
| --- | --- | --- | --- | --- | --- | --- | --- |
| CHROM | COMMON | NW&NE | CI | NPG | SPG | SSG | Total |
| HiC_scaffold_1 | 29302 | 4953 | 2328 | 1420 | 1155 | 739 | 39897 |
| HiC_scaffold_2 | 36246 | 5203 | 3359 | 1301 | 706 | 836 | 47651 |
| HiC_scaffold_3 | 23192 | 3474 | 1559 | 959 | 614 | 523 | 30321 |
| HiC_scaffold_4 | 25061 | 3059 | 1304 | 1103 | 1151 | 834 | 32512 |
| HiC_scaffold_5 | 25852 | 4350 | 2202 | 1670 | 878 | 912 | 35864 |
| HiC_scaffold_6 | 19792 | 3367 | 1040 | 561 | 597 | 1165 | 26522 |
| HiC_scaffold_7 | 60885 | 7240 | 3643 | 3688 | 1975 | 1490 | 78921 |
| HiC_scaffold_8 | 25575 | 4373 | 1568 | 1285 | 1069 | 736 | 34606 |
| HiC_scaffold_9 | 13875 | 2231 | 802 | 365 | 448 | 511 | 18232 |
| HiC_scaffold_10 | 20103 | 2629 | 1593 | 641 | 606 | 952 | 26524 |
| HiC_scaffold_11 | 22408 | 3082 | 1513 | 837 | 1198 | 493 | 29531 |
| HiC_scaffold_12 | 20036 | 2572 | 1068 | 1007 | 1131 | 1141 | 26955 |
| HiC_scaffold_13 | 57877 | 8118 | 4746 | 3423 | 1809 | 1697 | 77670 |
| HiC_scaffold_14 | 39204 | 4457 | 2026 | 1217 | 1282 | 513 | 48699 |
| HiC_scaffold_15 | 28636 | 3239 | 2725 | 1455 | 989 | 947 | 37991 |
| HiC_scaffold_16 | 28805 | 4836 | 2135 | 1295 | 1045 | 862 | 38978 |
| HiC_scaffold_17 | 10091 | 917 | 824 | 652 | 408 | 358 | 13250 |
| HiC_scaffold_18 | 23564 | 4150 | 2410 | 1011 | 751 | 791 | 32677 |
| HiC_scaffold_19 | 23045 | 3498 | 951 | 974 | 839 | 823 | 30130 |
| HiC_scaffold_20 | 21409 | 2675 | 1647 | 973 | 493 | 551 | 27748 |
| HiC_scaffold_21 | 22778 | 3363 | 1782 | 1498 | 802 | 439 | 30662 |
| HiC_scaffold_22 | 19092 | 3400 | 1750 | 1693 | 1100 | 396 | 27431 |
| HiC_scaffold_23 | 35499 | 4410 | 2024 | 1790 | 908 | 889 | 45520 |
| HiC_scaffold_24 | 65244 | 7139 | 5007 | 3341 | 2111 | 1929 | 84771 |
| HiC_scaffold_25 | 32802 | 5060 | 3196 | 1878 | 968 | 605 | 44509 |
| HiC_scaffold_26 | 51088 | 6907 | 3616 | 1964 | 1249 | 1347 | 66171 |
| HiC_scaffold_27 | 44724 | 6622 | 3306 | 2406 | 1636 | 971 | 59665 |
| Genome-wide | 826185 | 115324 | 60124 | 40407 | 27918 | 23450 | 1093408 |
| proportion of total | 0.755605 | 0.105472 | 0.054988 | 0.036955 | 0.025533 | 0.021447 | 1 |

| Supplementary Table 7: Estimated population sizes of Asian elephants in the five management units in India as recorded in 2017 (EPE 2017) | | |
| --- | --- | --- |
| Region/population unit | Number of elephants | Total population |
| Northwestern+Northeastern |  | 12,224 |
| a.    Northwestern | 2,085 |  |
| b.    Northeastern | 10,139 |  |
| Central |  | 3,128 |
| Southern |  | 10,747 |
| a.    N of Palghat Gap | 9,255 |  |
| b.    Between PG and SG | 1,327 |  |
| c.     South of Shencottah Gap | 165 |  |
| Reference: EPE (2017) Synchronized elephant population estimation India 2017. Project Elephant Division, Ministry of Environment, Forest and Climate Change, New Delhi. | | |

Supplementary Table 8: Model comparisons for fastsimcoal simulations. “agriculture” models use 7,000ya as the upper limit for divergence time and the “old_div” models use 17,500ya as the minimum divergence time. v2 and v3 refer to models presented in figure 7a and 7b respectively.

| **model** | **log likelihood** | **no. of parameters** | **AIC** | **delta** | **exp(-0.5delta)** | **weight** |
| --- | --- | --- | --- | --- | --- | --- |
| agriculture_v2 | -1721865.1 | 9 | 3443748.26 | 8859.098 | 0 | 0 |
| old_div_v2 | -1717607.3 | 9 | 3435232.64 | 343.482 | 2.5932E-75 | 2.5932E-75 |
| agriculture_v3 | -1721601.5 | 9 | 3443220.97 | 8331.812 | 0 | 0 |
| old_div_v3 | -1717435.6 | 9 | 3434889.16 | 0 | 1 | 1 |

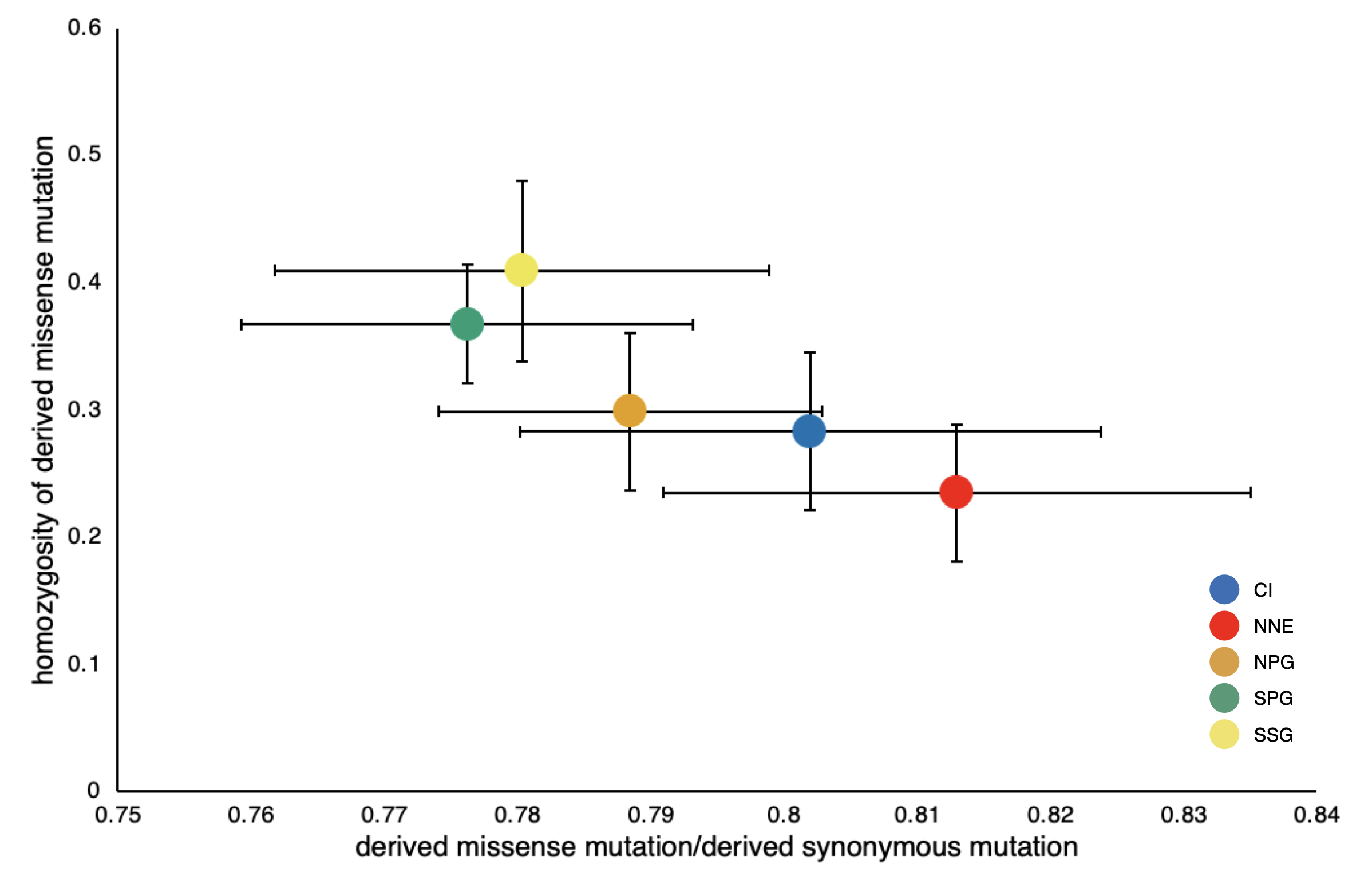
 Supplementary Figure 8: Mutation load measured as a function of homozygosity vs number of derived missense mutations. The number of derived deleterious alleles/number of derived neutral alleles is a proxy for number of deleterious alleles. The error bars indicate standard deviations.

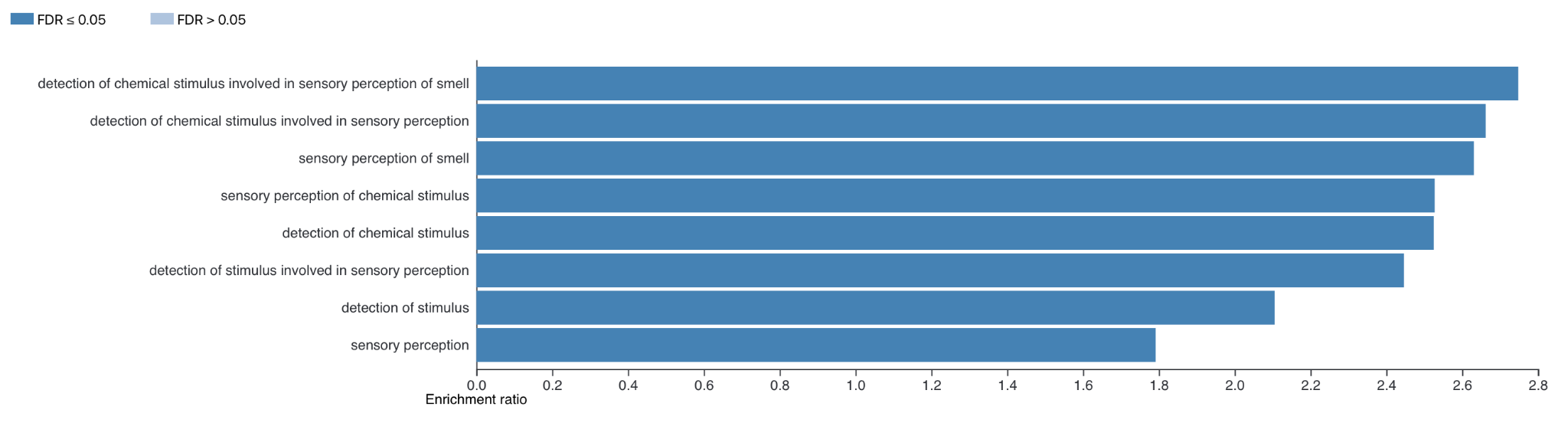

Supplementary Figure 9: Biological process affected by genes affected with missense mutations

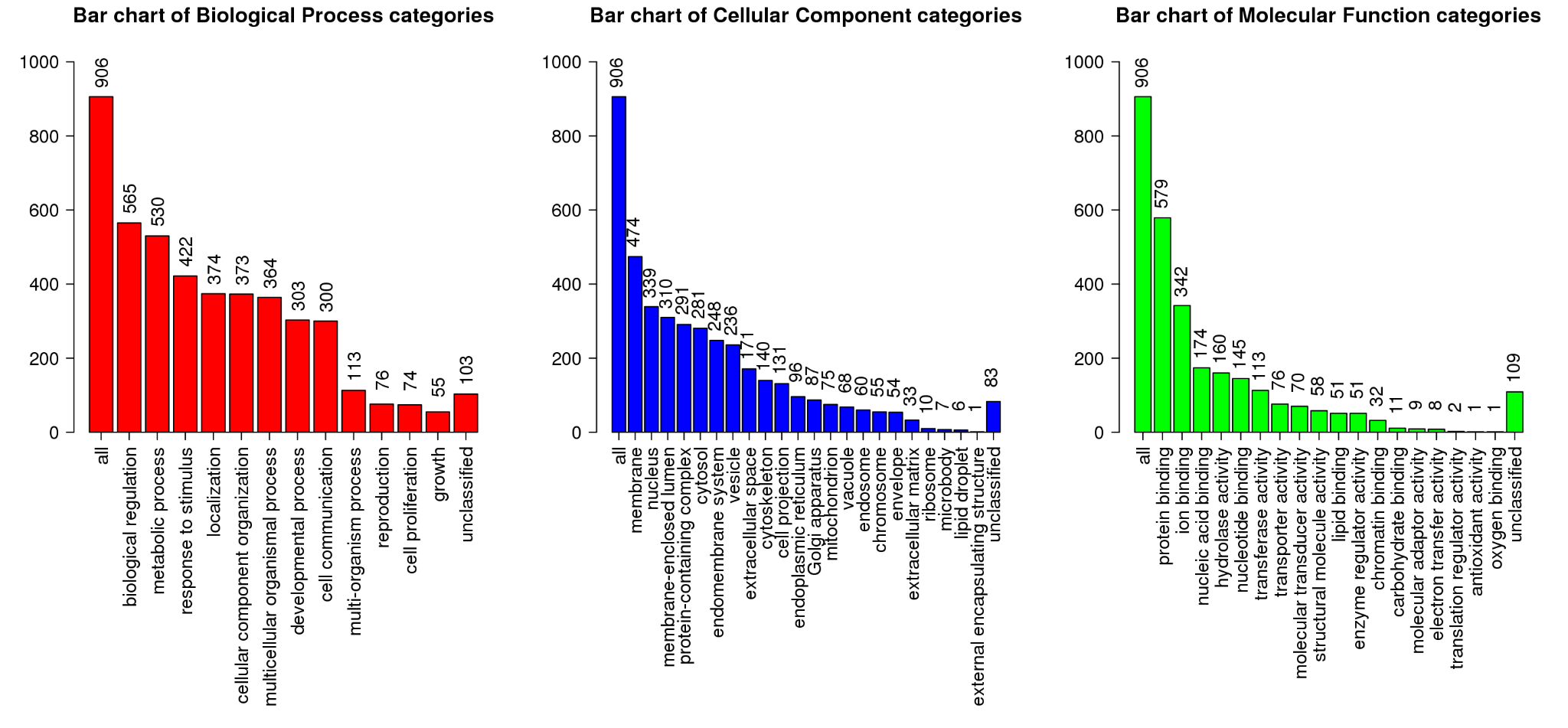

Supplementary Figure 10: Functions of genes with loss-of-function mutations

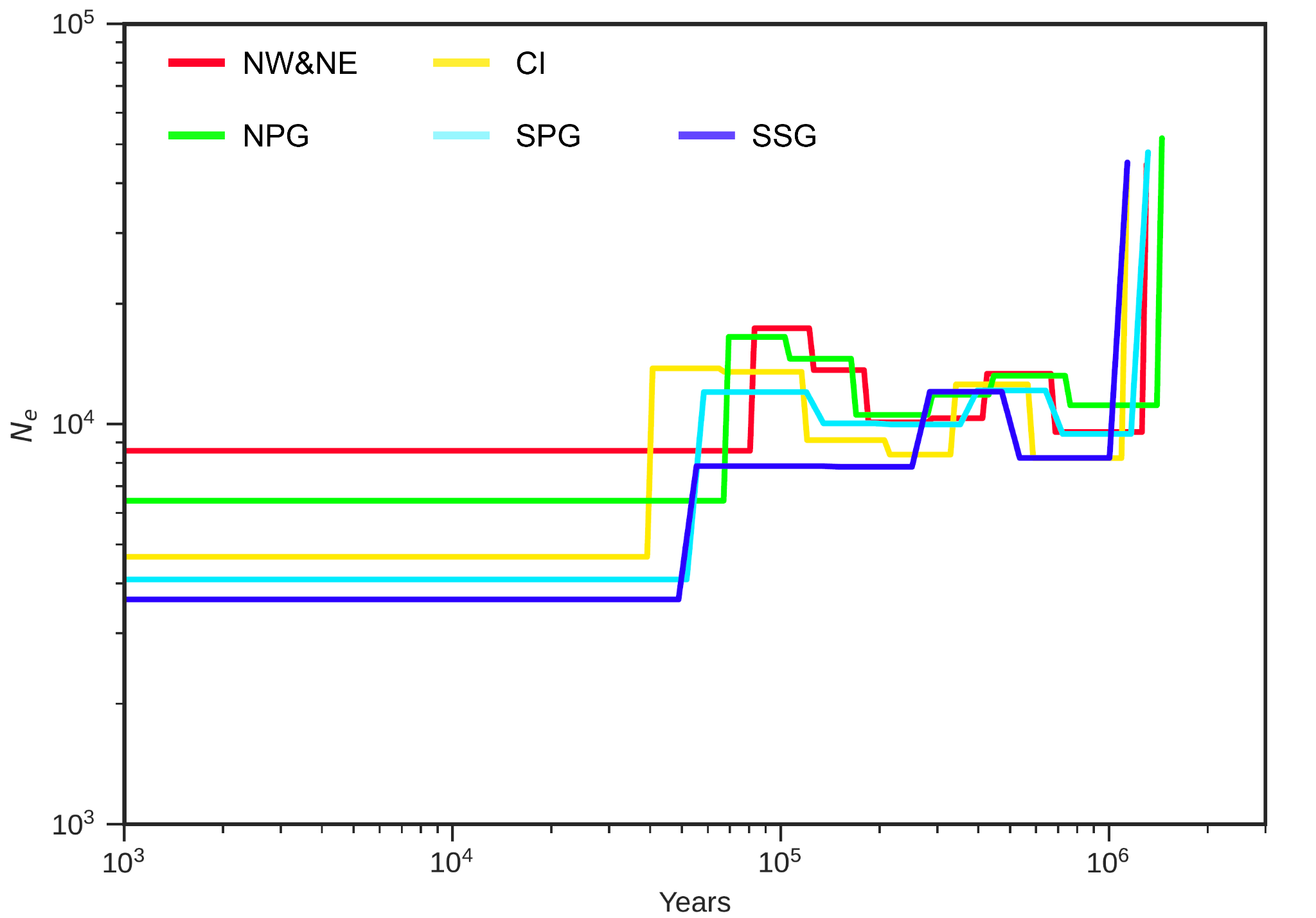

Supplementary Figure 11: Demographic history estimated by SMC++

EPE 2017. Synchronized elephant population estimation India 2017, Project Elephant Division, Ministry of Environment and Forests, New Delhi

Garcia‐Erill, G. and Albrechtsen, A., 2020. Evaluation of model fit of inferred admixture proportions. *Molecular Ecology Resources*, *20*(4), pp.936-949.

Leigh, JW, Bryant D (2015). PopART: Full-feature software for haplotype network construction. *Methods in Ecology and Evolution,* 6(9):pp.1110–1116.

Reddy, P.C., Sinha, I., Kelkar, A., Habib, F., Pradhan, S.J., Sukumar, R. and Galande, S., 2015. Comparative sequence analyses of genome and transcriptome reveal novel transcripts and variants in the Asian elephant Elephas maximus. *Journal of Biosciences*, *40*, pp.891-907.

Swofford, D. L. 2003. PAUP*. Phylogenetic Analysis Using Parsimony (*and Other Methods). Version 4. Sinauer Associates, Sunderland, Massachusetts.
